# High-Throughput Morphological Profiling of Patient-Derived Human Liver Organoids for Hepatotoxicity Hazard Identification and Compound Prioritization

**DOI:** 10.64898/2026.09.29.755355

**Authors:** Sophia R. Meyer, Max A. Garcia, Neil Zhao, Samantha F. Boisvert, Matthew McConnachie, Charles J. Zhang, Benjamin S. Halligan, Andrew M. Tidball, Andrew Tucker, Josepha Kaganove, Jesse W. Wotring, Joel A. Yates, Diana Flores, Robert J. Fontana, Jonathan Z. Sexton

**Author notes:** Indicates equal contribution (Sophia R. Meyer, Max A. Garcia, Neil Zhao).

## Abstract

Drug-induced liver injury (DILI) remains a major challenge in drug development because existing in vitro models poorly predict clinical hepatotoxicity. We coupled patient-derived human liver organoids (HLOs) with high-content imaging and automated machine learning (ML) to detect hepatotoxicity-associated phenotypes by morphological profiling. HLOs from six DILI patients and one healthy donor (n = 7) retained donor-specific baseline phenotypes in hepatocyte maturity, metabolism, and morphology. ML models trained per line on MAD-robustized, well-level-aggregated median features achieved compound-level AUROCs of 0.88 (95% CI 0.86-0.90) to 0.91 (95% CI 0.89-0.93) and balanced accuracies of 0.81-0.84 in distinguishing 54 hepatotoxicants from 20 non-hepatotoxic controls. Averaging predictions across all seven lines at an optimized threshold gave 94.4% sensitivity (95% CI 84.6-98.8%) and 85.0% specificity (95% CI 62.1-96.8%), substantially outperforming HLO cytotoxicity assays and cancer cell line profiling. Lines were moderately-to-highly concordant (Pearson r = 0.66-0.85), and benchmark concentration modeling revealed reproducible inter-line differences in phenotypic sensitivity (median 4.0-fold, several compounds exceeding 20-fold). Exposure anchoring of benchmark concentrations to human-equivalent plasma concentrations, using in vitro mass-balance dosimetry and curated plasma free fractions, placed organoid points of departure within 100-fold of clinical C_max_ for 60 of 66 compounds (23-fold residual offset, 95% CI 12-45), and placed two of the three misclassified negative controls an order of magnitude above clinical C_max_. Sulindac injury signatures were significantly enhanced by TNFα co-treatment, demonstrating capacity to detect pro-inflammatory-mediated DILI. This platform preserves inter-line variability while enabling reproducible, scalable multi-line consensus hazard classification; because compounds were tested over a common nominal concentration range, the readout defines hepatocellular hazard, which must be combined with exposure as an independent input to estimate clinical DILI risk.

## Introduction

The FDA’s recent emphasis on qualifying human preclinical models and other New Approach Methodologies (NAMs) highlights a critical need for translational in vitro systems derived from human cells and tissues to improve the predictive accuracy of preclinical toxicology.^1–3^ Among adverse drug reactions, drug-induced liver injury (DILI) remains a major challenge in drug development and is a leading cause of clinical trial failure and post-marketing withdrawal.^4–6^ Traditional toxicology approaches, including animal models and primary human hepatocyte monocultures, often fail to capture human-specific liabilities, limiting their utility for early safety de-risking. As pressure mounts to "fail fast and fail cheap," there is a growing need for scalable platforms capable of predicting patient-relevant toxicities earlier in the drug development pipeline.

The complex architecture of the liver and its multicellular interactions are central to many DILI phenotypes.^7^ Hepatocytes carry out most xenobiotic metabolism, while non-parenchymal cells modulate injury outcomes: Kupffer cells amplify inflammatory signaling,^8^ and hepatic stellate cells mediate fibrotic responses.^9^ These multicellular dynamics are absent in conventional 2D hepatocyte monocultures, limiting their physiological relevance. Moreover, standard hepatotoxicity screening relies heavily on cytotoxicity-based readouts, which can miss early or non-lethal phenotypic changes associated with hepatocellular stress.

NAMs aim to address these limitations by enabling human-relevant, mechanistically informative, and scalable systems for toxicity assessment. Patient-derived human liver organoids (HLOs) are 3D in vitro microtissues generated from induced pluripotent stem cells that recapitulate some key liver cell types, including hepatocytes, hepatic stellate cells, and Kupffer cells.^10,11^ Our prior characterization demonstrated reliable production of functional and metabolically-competent hepatocyte-like cells, hepatic stellate cells with a minor population (∼1%) of Kupffer-like macrophages.^12,13^ Functional assessment showed significant CYP1A, CYP2B, and CYP3A metabolic activity as measured by substrate turnover, albumin secretion in dispersed monolayers of ∼5 µg/day/10^6^ cells for 7 days, and sensitivity to known hepatotoxicants over multi-week culture periods.^12,13^ Additionally, adapting HLOs to MPS systems significantly enhanced the functional duration to 14-28 days.^13^ These systems have shown promise for modeling drug metabolism, hepatic injury, and inter-individual variability in drug responses.^12–14^ In parallel, high-content morphological profiling coupled with ML has emerged as a powerful approach for capturing complex cellular responses to chemical perturbations, outperforming traditional cytotoxicity-based methods in toxicology and drug safety applications.^15,16^

In this study, we present a high-throughput NAM that integrates patient-derived HLOs with multiplexed high-content imaging and an automated machine-learning pipeline to detect hepatotoxicity-associated phenotypes. Using a panel of 74 compounds, including 54 with established human DILI risk and 20 without clinically apparent hepatotoxicity, we evaluated model performance across HLOs and immortalized cell lines. HLOs were dispersed into 384-well plates and exposed to drugs across five concentrations for 72 hours. High-content multiplexed fluorescent imaging targeting mitochondrial activity, lipid accumulation, nuclear morphology, and cellular architecture was used to identify phenotypic injury signatures associated with hepatotoxicity phenotypes. We assessed hepatotoxicity hazard for each drug using morphological profiling-based ML classification in HLOs derived from six donors who experienced DILI, one healthy control, and in four cancer cell lines (HepG2, HUH7, U2OS, and HCT116). This platform provides a scalable framework for high-throughput hepatotoxicity hazard identification and mechanistic hypothesis generation that can be used in early-stage drug development and can inform medicinal chemistry optimization during hit-to-lead and lead optimization. We position the assay deliberately as a first-tier hazard identification and compound-ranking tool rather than as a predictor of clinical DILI risk for an individual candidate. Because compounds were tested over a common nominal concentration range irrespective of therapeutic exposure, protein binding, or hepatic accumulation, the readout defines hepatocellular hazard.

This work extends previous organoid-based hepatotoxicity screening in four respects. First, the donor panel is derived from patients with adjudicated DILI Network causality scores, a resource that cannot be reconstituted from commercially available lines. Second, we show that consensus predictions across genetically distinct lines outperform any single line, and that pairwise concordance identifies which lines carry complementary rather than redundant information, providing a design principle for constructing a donor panel. Third, feature-level benchmark concentration modeling converts the platform from a binary classifier into a tool for ranking compounds and quantifying variability between lines. Fourth, TNFα co-treatment prospectively recovers a compound missed under basal screening conditions, with the sulindac sulfide metabolite arm establishing the mechanistic basis. Relative to conventional single-line, viability-anchored screens, the present study establishes a multi-donor panel assessed by morphological profiling with concentration-response points of departure with high predictive accuracy.

## Results

### Generation and Phenotypic Characterization of Patient-Derived Human Liver Organoids from Seven Donors

To assess donor-to-donor variability and capture genetic diversity within the DILI cohort, we generated six patient-derived induced pluripotent stem cell (iPSC) lines from donors enrolled in the DILI Network (DILIN) with high single-drug causality scores **(Supplemental Table S1)**.^17^ Each iPSC line was reprogrammed from expanded peripheral blood erythroid progenitors using episomal reprogramming, a well-established non-integrating method for generating iPSCs from blood samples.^17^ As a non-DILI control, we included the well-characterized iPSC line 72.3, derived from foreskin fibroblasts of a healthy donor with no history of DILI.^18^ Together, these lines comprise a panel of seven iPSC models (six DILI, one non-DILI control) used throughout this study.

Single nucleotide polymorphism (SNP) DNA microarray analysis confirmed the identity and genomic integrity of all iPSC lines, with the exception of line 71G, which exhibited a ∼0.5 Mb duplication at chr7:33,077,502–33,579,232 encompassing most of the BBS9 gene and may represent a potential confounding factor when interpreting its drug-response phenotype.

All seven iPSC lines were differentiated into human liver organoids (HLOs) using a previously established protocol.^10,11,13^ Hereafter, we refer to organoids derived from each iPSC line as a distinct HLO model, reflecting donor-specific genetic and phenotypic backgrounds. The resulting HLO models were dissociated into a single cell suspension, seeded into 384-well plates, and treated with a curated library of 74 compounds, including 54 hepatotoxic (positive controls, PC) and 20 non-hepatotoxic drugs (negative controls, NC) (**Supplemental Table S2**). Compounds were tested at five concentrations in triplicate, distributed across three 384-well plates with 32 DMSO vehicle control wells per plate. This yielded 1,110 compound-treated and 42 DMSO controls wells per HLO model. After 72 hours of exposure, cells were fixed, stained, and imaged, followed by single-cell phenotypic profiling (**Figure 1A**). **Figure 1B** shows representative images of the fluorescent probe set, shown from left to right; DNA/lipids/mito, DNA/HNF4α, and HCS CellMask Orange (note: 100 μm scale bar, uniform scaling). High-content imaging coupled with segmentation and feature extraction enabled quantification of key cellular features, including nuclear morphology, mitochondrial activity, lipid accumulation, and cytoplasmic architecture. These features provided readouts including hepatocyte identity (HNF4α), steatosis, oxidative stress, and overt cytotoxicity phenotypes (**Figure 1C**).

**Figure 1.**
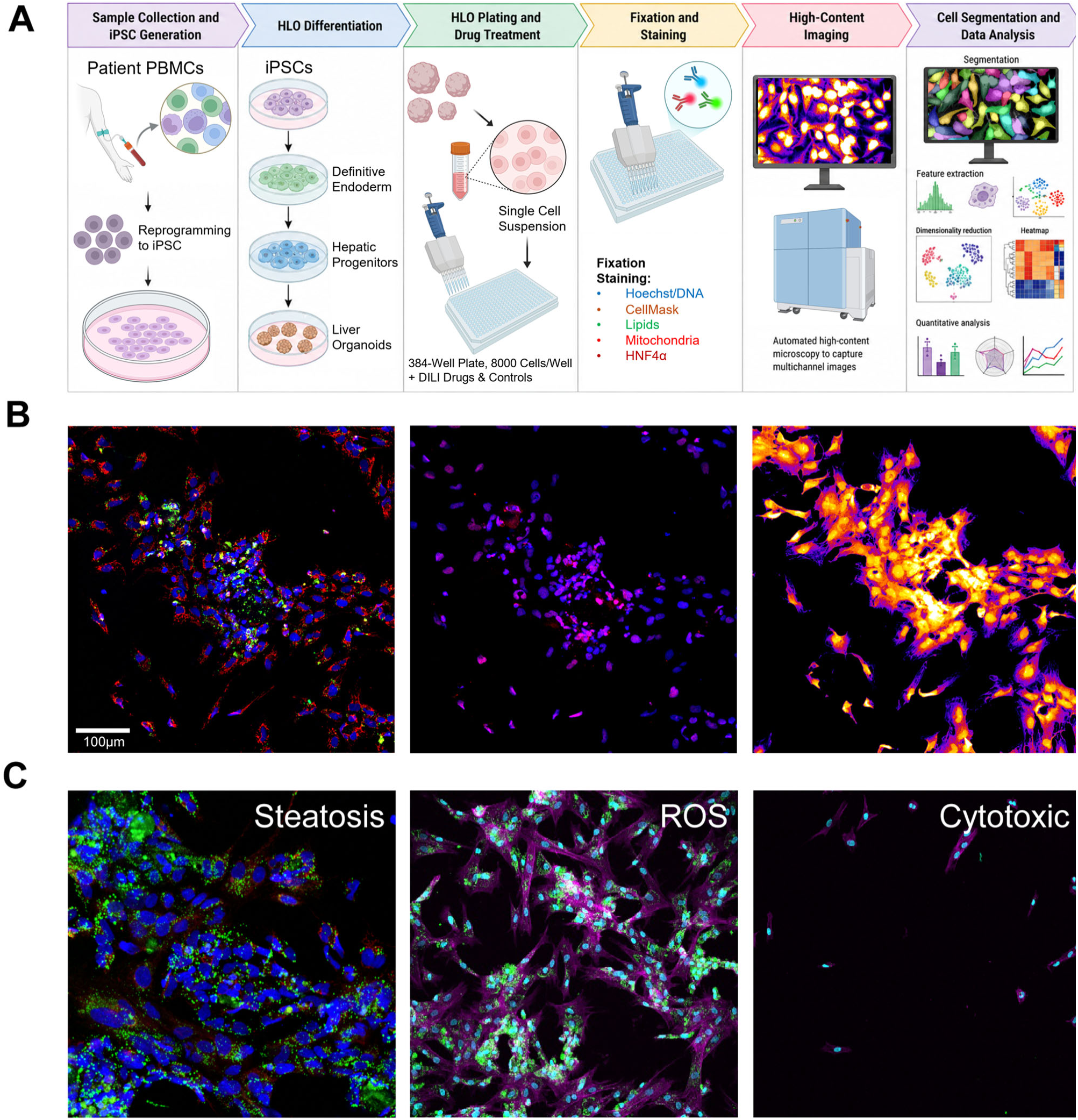
High-throughput DILI risk prediction platform in human liver organoids (HLOs). **(A)** Workflow schematic. Patient blood samples are reprogrammed into iPSCs and differentiated into HLOs containing predominantly hepatocyte- and stellate-like cells. HLOs are dissociated, seeded into multi-well plates, and treated with test compounds for 72 h. Cells are then fixed, stained with fluorescent markers, and imaged by high-content fluorescence microscopy. Images are computationally segmented and features extracted for machine-learning–based analysis. **(B)** Representative images of cellular composition and subcellular markers. *Left:* Hoechst (blue), neutral lipid droplets (green), mitotracker (red). *Middle:* Hoechst (blue), HNF4α (red, hepatocyte-like cells). *Right:* HCS CellMask Orange. Scale bar = 100 µm. **(C)** Representative DILI-relevant toxicity phenotypes induced by reference compounds: steatosis, oxidative stress (ROS), and cytotoxicity.

Pre-treatment phenotypic characterization revealed both shared and distinct baseline features across the seven HLO models. Uniform Manifold Approximation and Projection (UMAP) analysis showed model-specific phenotypic distributions with partial overlap (**Figure 2A**), consistent with donor-dependent variation within a conserved phenotypic framework. Pairwise Wasserstein distance analysis of vehicle-treated controls further quantified these relationships, demonstrating both separation and relative similarity among HLO models (**Figure 2B**). For example, lines 63D, 65C, and 77A formed the most phenotypically similar cluster, while 64B, 71G, and 83C showed intermediate similarity to each other. The healthy donor line 72.3 was the most phenotypically distinct.

**Figure 2.**
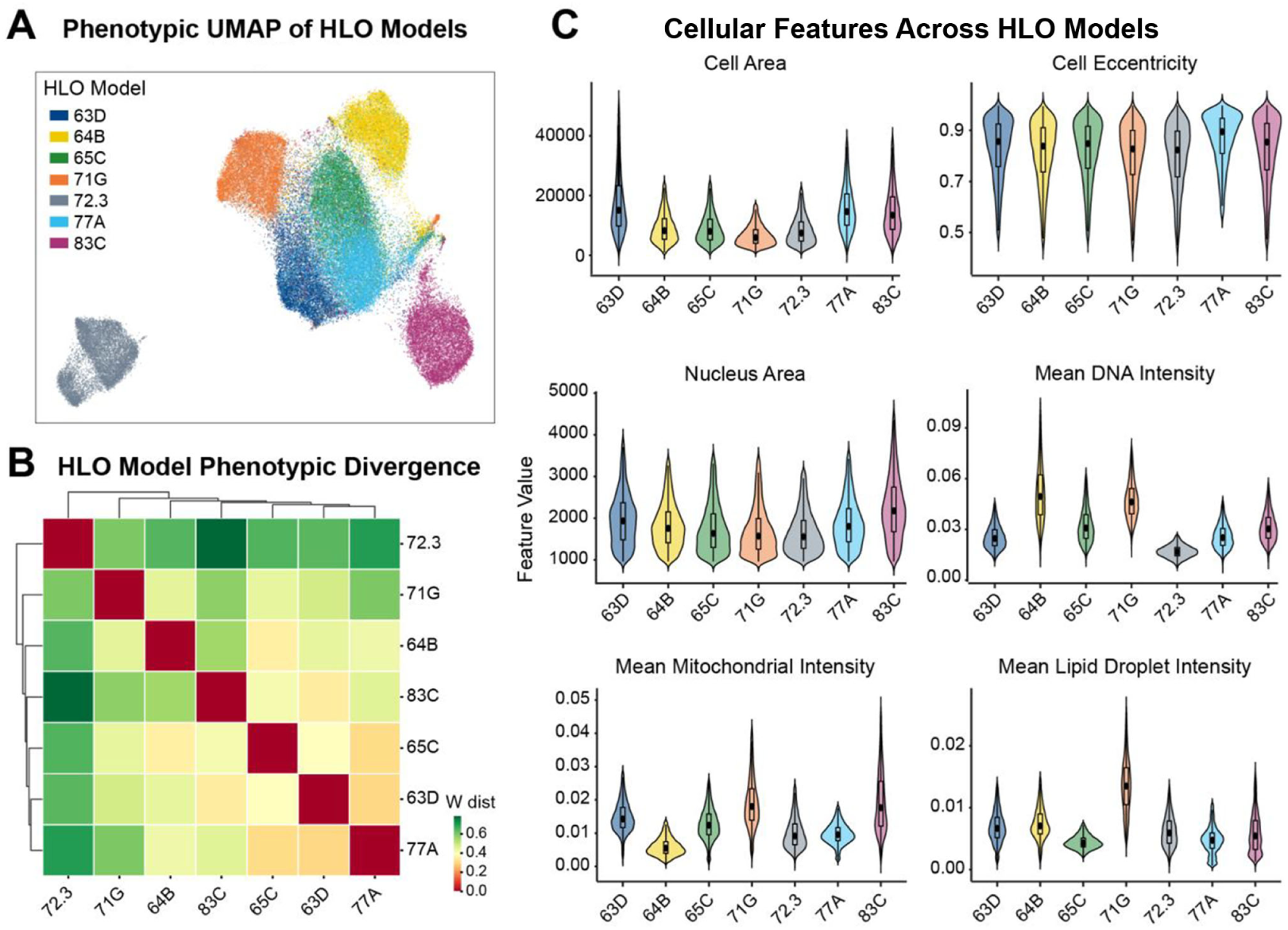
Baseline phenotypic heterogeneity across human liver organoid (HLO) donor models. **(A)** UMAP projection of single-cell phenotypes from vehicle-treated, dispersed HLOs (n = 10,000 cells per model; nn = 30, md = 0.1; silhouette = 0.378). Each point represents one cell, colored by donor model. **(B)** Pairwise Wasserstein distances between donor models computed on single-cell phenotypic feature vectors; lower values indicate greater similarity. **(C)** Violin plots of representative cellular features across the seven HLO models: cell area, cell eccentricity, nucleus area, and mean DNA, mitochondrial, and lipid droplet intensities. Boxes and center points denote interquartile range and median.

Feature-level comparisons supported this balance between variability and conservation (**Figure 2C**). While specific differences were observed, such as elevated lipid droplet intensity in 71G and lower mitochondrial intensity in 64B, many core cellular features showed overlapping distributions across models. Statistical testing and effect size analysis confirmed that, despite significant inter-donor differences, these phenotypes exist within a shared morphological framework (**Supplemental Figure S1A, B**). Together, these results indicate that HLOs retain donor-specific phenotypic signatures while preserving a common baseline architecture, providing a consistent yet biologically diverse platform for downstream drug-response studies.

### Machine Learning on Morphological Profiles Discriminates Hepatotoxicants at the Compound Level

DILI risk prediction ML models were trained independently on HLOs derived from each donor using AutoGluon AutoML with compound-grouped five-fold cross-validation, where all wells from a given compound were assigned exclusively to either training or validation folds to prevent data leakage across concentration replicates. Out-of-fold (OOF) predictions were generated for each well using ensemble models trained only on held-out compounds, ensuring unbiased performance estimates.

Across each donor-derived HLO population, predicted DILI probabilities demonstrated a clear separation between hepatotoxic and non-hepatotoxic compounds at the well level (**Figure 3A**). Classification performance was consistently high across donors, with AUROC values ranging from 0.88 (95% CI 0.86-0.90) to 0.91 (95% CI: 0.89–0.93) (**Figure 3B**) and area under the precision-recall curves (AUPRC) of approximately 0.95 across models (**Figure 3C**). Balanced accuracy ranged from 0.81 to 0.84 (mean = 0.83), with Matthews correlation coefficients (MCC) ranging from 0.59 to 0.65 (mean = 0.63), indicating robust and well-calibrated classification across donors (**Supplemental Figure S2A**). Notably, the healthy donor line 72.3 achieved performance comparable to the best-performing DILI patient-derived lines. The highest-performing individual model was derived from donor line 64B (AUROC 0.91, AUPRC 0.96, balanced accuracy 0.84). These results indicate that morphological features captured by multiplexed imaging encode strong predictive signals of hepatotoxicity across diverse genetic backgrounds.

**Figure 3.**
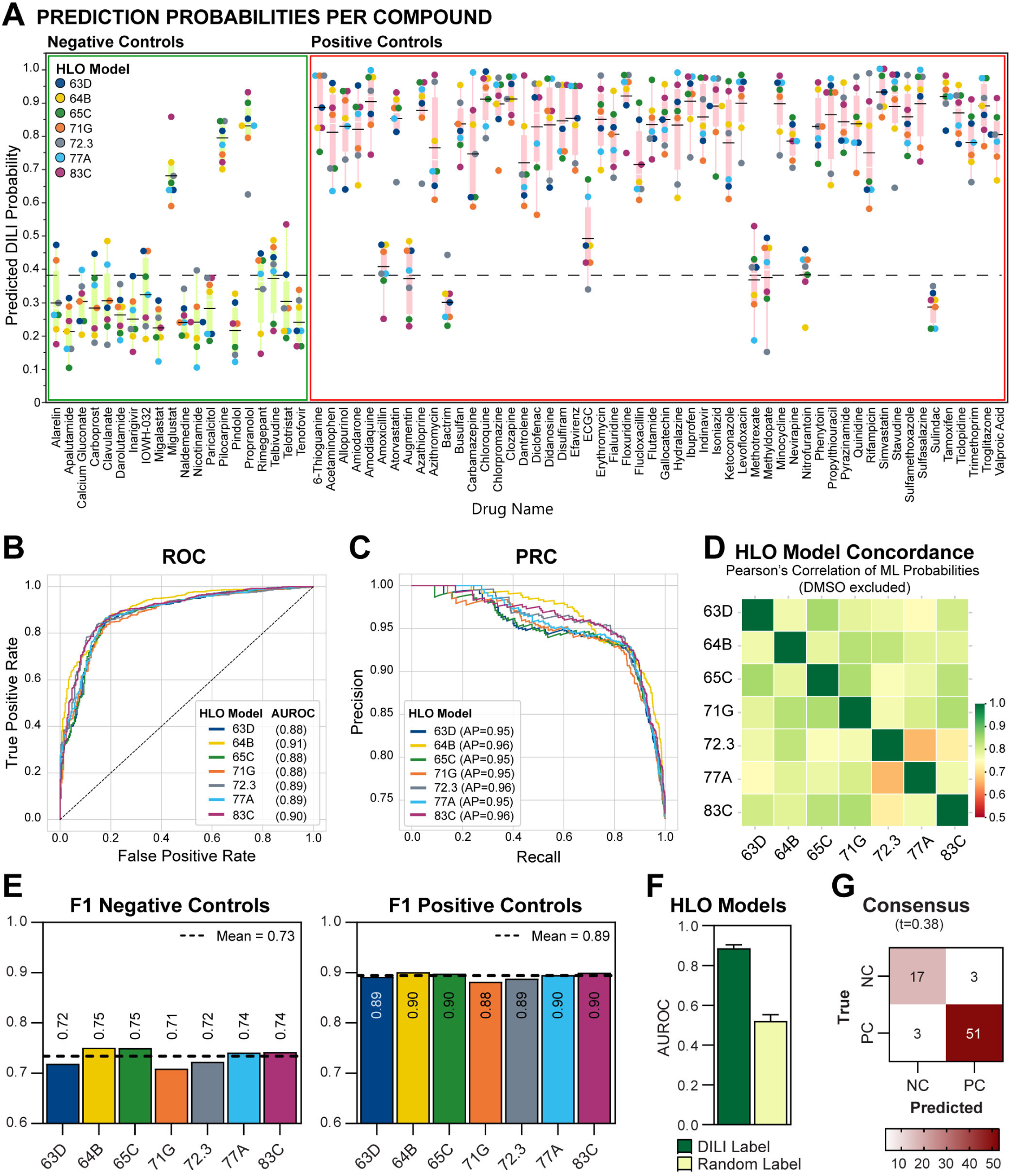
Phenotypic machine learning across HLO models enables robust and generalizable prediction of DILI risk. **(A)** Out-of-fold (OOF) predicted DILI probabilities for 74 compounds across the seven donor-derived HLO models, grouped by ground-truth negative (left) and positive (right) controls. Each point represents the prediction from one HLO model; the dashed line indicates the classification threshold = 0.38; the black line indicates the consensus prediction score. **(B)** Receiver operating characteristic (ROC) curves per HLO model; AUROC values shown in the legend. **(C)** Precision–recall curves per HLO model; average precision (AP) values shown in the legend. **(D)** Pairwise Pearson correlation matrix of compound-level OOF predicted probabilities across HLO models (DMSO excluded). **(E)** Per-model F1 scores for negative (left) and positive (right) controls; dashed lines indicate the cross-donor mean. **(F)** AUROC under true versus label-permuted DILI labels (mean ± SD); permutation reduces performance toward chance, indicating absence of substantial label leakage. **(G)** Confusion matrix for the consensus model (mean prediction across HLO models, threshold = 0.38).

To assess whether different donor-derived HLO models produce related drug-response predictions, we computed pairwise Pearson correlations of OOF predicted probabilities that showed moderate-to-high correlation across donors (**Figure 3D**), indicating shared predictive structure while preserving model-specific variation. Correlation coefficients ranged from r = 0.66 (72.3 vs 77A; most discordant) to r = 0.85 (63D vs 65C; most concordant). This pattern suggests that some HLO models contribute overlapping information, whereas more divergent models provide complementary predictive coverage. Consistent with this observation, integrating predictions across donors improved classification performance relative to any single patient-derived HLO (see **Figure 3G**; per-line comparison, **Supplemental Figure S2B, D**).

Model interpretability was supported by performance on control compounds. Classifier performance was evaluated using the F1 score, the harmonic mean of precision and recall. F1 scores for NCs were consistently above 0.7 (mean = 0.73), while F1 scores for PC drugs were even higher (mean = 0.89) across models (**Figure 3E**), indicating robust discrimination of non-hepatotoxic and hepatotoxic compounds across all HLO models. These trends were consistent with confusion matrices, which showed strong true positive and true negative classification across both individual and consensus HLO models (**Figure 3G**, **Supplemental Figure S2D**).

To confirm that classifier performance reflected genuine biological signal rather than artifacts of model training (overfitting), we performed a label-permutation control: class labels were randomly shuffled and the full training and evaluation pipeline was repeated. Under permutation, AUROC values approached 0.5 (**Figure 3F**), consistent with random classification and supporting the validity of the modeling framework.

A consensus DILI prediction was derived by averaging the predicted probability across all seven HLOs per compound and thresholding at the Youden-J optimal cutoff of 0.38. This rule correctly identified 51 of 54 hepatotoxicants (sensitivity 94.4%, 95% CI 84.6%–98.8%) and 17 of 20 non-hepatotoxicants (specificity 85.0%, 95% CI 62.1%–96.8%), yielding a balanced accuracy of 89.7% (**Figure 3G**). The consensus substantially outperformed any individual HLO model (per-line balanced accuracy 0.81–0.84; **Supplemental Figure S2B, D**) and the cytotoxicity-based approach described below. Because the threshold was optimized on the same 74-compound panel used for evaluation, these are internally optimized performance estimates and will require validation in an independent compound set to confirm generalizability. Together, these results demonstrate that phenotypic ML applied to multi-donor HLO morphological profiles enables robust compound-level discrimination of hepatotoxicants from non-hepatotoxic controls, and that aggregating responses across genetically diverse donors substantially improves classification performance. These estimates describe detection of hepatotoxicity-associated phenotypes under the tested concentration range and are not, on their own, estimates of clinical DILI risk.

### Morphological Profiling Coupled with Machine Learning Outperforms Cytotoxicity in Hepatotoxicity Assessment

To assess the combined impact of model system and analytical approach on hepatotoxicity prediction, we compared ML-based morphological profiling and viability-based classification across the seven patient-derived HLOs and four immortalized cell lines: HepG2 and HUH7 (hepatocellular carcinoma), HCT116 (colorectal carcinoma), and U2OS (osteosarcoma). Cytotoxicity was scored from cell counts obtained on the same assay plates used for morphological profiling (classification criteria in **Figure 4** legend).

**Figure 4.**
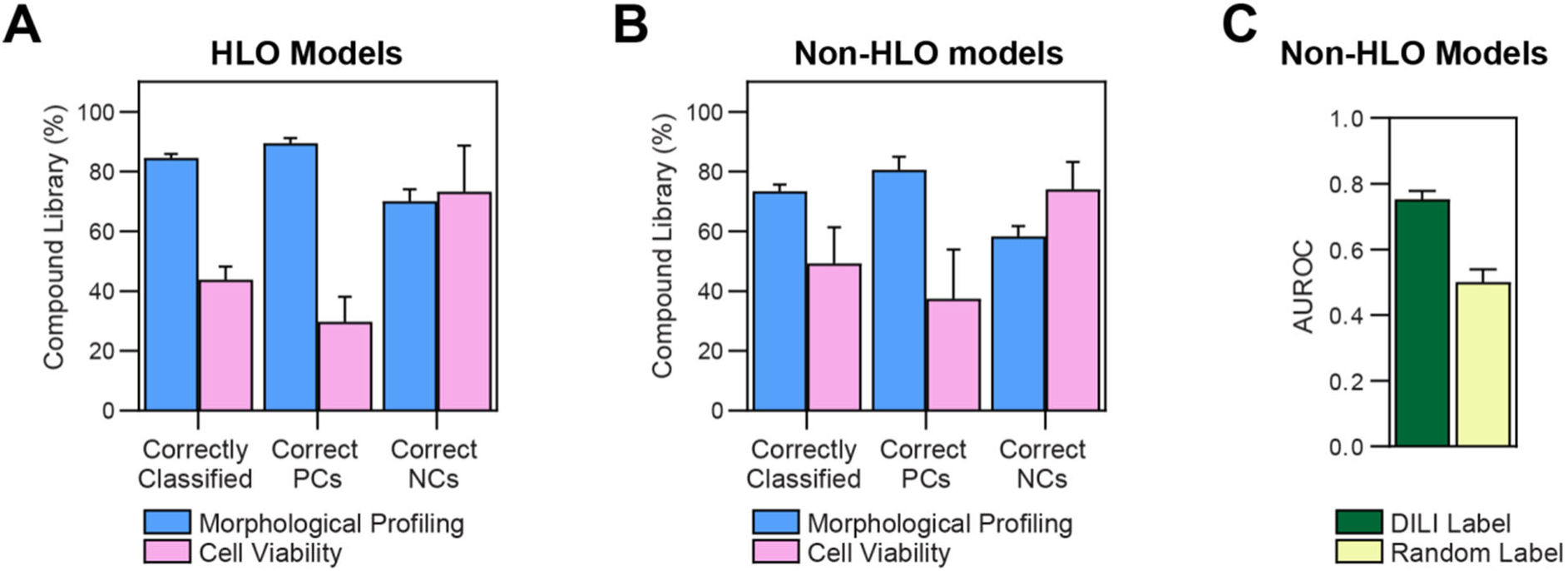
Morphological profiling outperforms viability-based classification for DILI prediction across HLO and non-HLO models. **(A)** Percentage of the 74-compound library correctly classified across HLO models by machine learning on morphological features versus cell viability. For viability-based classification, positive controls (PCs) were scored correct if any tested concentration showed viability <50%; negative controls (NCs) required viability >50% at all tested concentrations. Bars show overall accuracy, correctly classified PCs, and correctly classified NCs (mean ± SD across HLO models). **(B)** As in (A), for non-HLO comparator cell models. **(C)** AUROC for non-HLO models under true versus label-permuted DILI labels (mean ± SD).

In HLO models, morphological profiling substantially outperformed viability-based assessment (**Figure 4A**). Across the full compound library, ML achieved an average accuracy of 84.6%, compared to 43.8% for viability (McNemar p<0.0001). This improvement was driven by markedly higher sensitivity for PC compounds (∼3-fold increase), while NC classification was comparable between methods (70.0% vs 73.2%, p=0.47), indicating that gains in sensitivity did not come at the expense of specificity. A small subset of compounds including Augmentin, Bactrim, and sulindac was misclassified by both methods. These compounds are associated with idiosyncratic or immune-mediated mechanisms of liver injury that are not recapitulated in this in vitro system (**Supplemental Table S3**).

Performance declined in immortalized cell lines (**Figure 4B**). While ML still outperformed viability (73.3% vs 49.3% overall accuracy, McNemar p < 0.0001), its accuracy fell ∼11 percentage points relative to HLOs. The ML performance decrease was modest for PCs (80.5% vs 89.4%, p=0.005) but more pronounced for NCs (58.3% vs 70.0%, p=0.087, n.s.). Viability-based NC classification, by contrast, was nearly identical across systems (74.0% in cell lines vs 73.2% in HLOs, p=0.47), and the overall accuracy gap was attributable to elevated ML false-positive rates in immortalized lines. The corresponding ML-versus-viability gap on NC compounds was 15.7 percentage points in cell lines but only 3.2 in HLOs (**Figure 4A–B**), consistent with elevated morphological false-positive rates in immortalized lines and likely reflecting altered baseline morphology and amplified non-specific responses at higher compound concentrations. Despite reduced specificity, ML predictions in non-HLO models remained well above chance. AUROC was 0.75 for true DILI labels versus 0.50 for permuted labels (**Figure 4C**), confirming that the classifier captures biologically meaningful hepatotoxicity signals even outside a physiologically relevant context.

Together, these results demonstrate that both the in vitro liver model and the analytical readout independently improve DILI prediction accuracy. Morphological profiling improves sensitivity for hepatotoxic compounds, while a disease-relevant HLO system preserves specificity. Combining the two yields an effective in vitro platform for hepatotoxicity prediction.

### Benchmark concentration modeling reveals inter-line variability in phenotypic concentration–response

While supervised ML provided robust DILI predictions, this approach does not directly estimate the concentration at which phenotypic perturbations emerge above baseline variability. To quantify the concentration dependence of morphological responses and assess inter-individual variability in toxicological sensitivity, we applied benchmark dose (BMD) modeling via BMDExpress3 to the same CellProfiler-derived morphological features used in ML DILI prediction for the 54 DILI-associated PCs. This analysis identifies feature-level points of departure (PoDs) by determining the BMD at which each morphological feature shows a significant dose-dependent deviation from the DMSO controls for each donor–compound combination. All statistical analyses and visualizations were restricted to BMD values within the empirically tested dose range (1–100 µM) to exclude extrapolated values beyond observed concentrations.

Omnibus testing (Kruskal-Wallis and Anderson-Darling, FDR < 0.05) identified significant donor effects for all 54 compounds. Representative compounds illustrate a spectrum of variability observed across the donor panel (**Figure 5A**).

**Figure 5.**
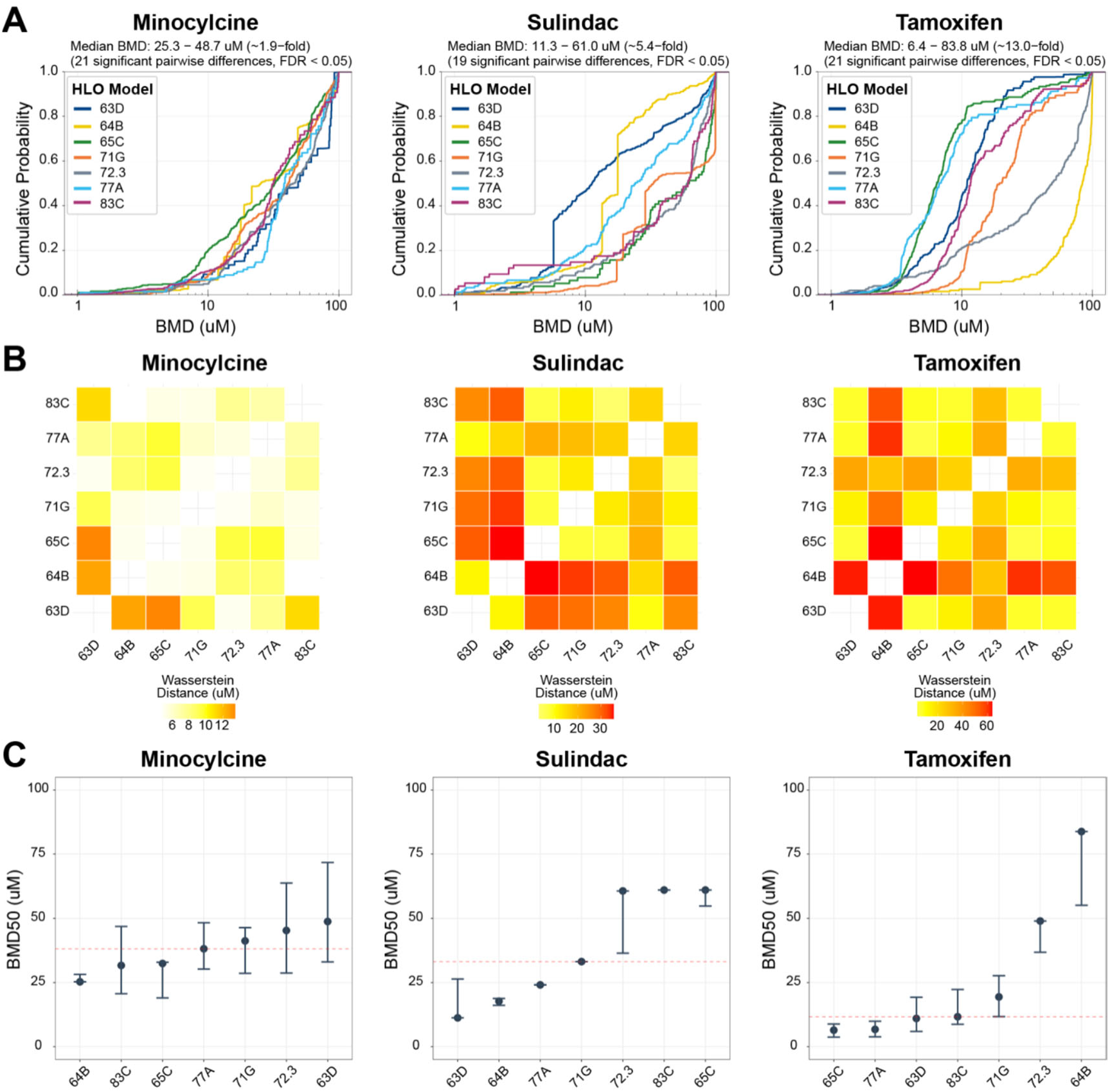
Inter-individual variability in benchmark dose distributions across HLO donor models. **(A)** Empirical cumulative distribution functions (eCDFs) of feature-level benchmark dose (BMD) values for three representative compounds spanning low (minocycline), intermediate (sulindac), and high (tamoxifen) inter-donor heterogeneity. Each curve represents one donor (n = 7). Median BMD ranges and fold differences across donors are indicated above each panel. Pairwise differences between donor distributions were assessed by Kolmogorov–Smirnov test with false discovery rate (FDR) correction. **(B)** Pairwise first-Wasserstein distance matrices (μM) of BMD distributions for the same three compounds; higher values indicate greater donor-to-donor divergence. **(C)** Median benchmark dose (BMD^50^) per donor for each compound. Error bars denote the BMDL^50^ (lower bound) and BMDU50 (upper bound) from BMDExpress3 best-fit dose–response models, and dashed red lines indicate the grand median BMD^50^ across donors. BMD, BMDL, and BMDU values were restricted to the tested dose range (≤100 μM) prior to analysis.

For minocycline, donor empirical cumulative distribution function (eCDF) curves largely overlapped across the tested dose range, consistent with a conserved phenotypic response profile and low inter-donor heterogeneity (donor median BMD range 25.3–48.7 µM; ∼1.9-fold; 21 significant pairwise differences, FDR < 0.05). In contrast, sulindac exhibited partial divergence among donor curves producing a biphasic distribution pattern consistent with intermediate heterogeneity (donor median BMD range 11.3–61.0 µM; ∼5.4-fold; 19 significant pairwise differences, FDR < 0.05). Tamoxifen displayed the highest variability, with donor curves broadly separated across the dose axis with minimal overlap, indicating high heterogeneity in phenotypic sensitivity (donor median BMD range 6.4–83.8 µM; ∼ 13.0-fold; 21 significant pairwise differences, FDR < 0.05).

Pairwise Wasserstein distances between donor BMD distributions quantified these differences on an absolute scale (**Figure 5B**). For minocycline, pairwise distances were generally low, with a maximum of 13.6 µM difference between a given donor pair. In contrast, sulindac exhibited distances up to 35.6 µM between the most divergent donor pairs, while tamoxifen reached a maximum pairwise distance of 63.8 µM, with 12 of 21 donor pairs exceeding the 10 µM threshold for meaningful differences. To complement the feature-level benchmark concentration (BMC) analysis, we compared the median benchmark dose (BMD_50_) and its 95% confidence bounds (BMDL_50_/BMDU_50_) from BMDExpress3 best-fit models for each compound/donor combination. These summary estimates recapitulated the inter-donor variability observed in the full feature-level distributions (**Figure 5C**). For tamoxifen, the BMDU_50_ of the most sensitive donor (65C) fell below the BMDL_50_ of the most resistant donor (64B), indicating non-overlapping confidence bounds across the donor range.

Global ranking of all 54 PC compounds by maximum pairwise Wasserstein distance revealed that the magnitude of inter-individual variability varied substantially across the compound panel and that all compounds exceeded the 10 µM effect size threshold (**Figure 6A**). Floxuridine and tamoxifen exhibited the greatest inter-donor BMD divergence (maximum pairwise distance > 60 µM), whereas compounds such as minocycline and allopurinol exhibited the least (< 15 µM). The number of significant pairwise comparisons per compound ranged from 9 to all 21 possible donor pairs. Hierarchical clustering of z-scored median BMD values (**Figure 6B**) revealed structure across both compound and donor dimensions. Donors 77A and 65C co-clustered and were separated from donors 72.3 and 64B by the primary column split, suggesting shared response phenotypes within subsets of the donor cohort. Across all compounds, the median fold variability between the most sensitive and most resistant donors was ∼4.0-fold, with several compounds exceeding ∼20-fold variability. At the global level, eCDFs of BMD values pooled across all compounds (**Figure 6C**) showed that donor-level separation persisted even after aggregation over the full compound panel, with donor median BMD values that ranged from 20.8 to 41.1 µM.

**Figure 6.**
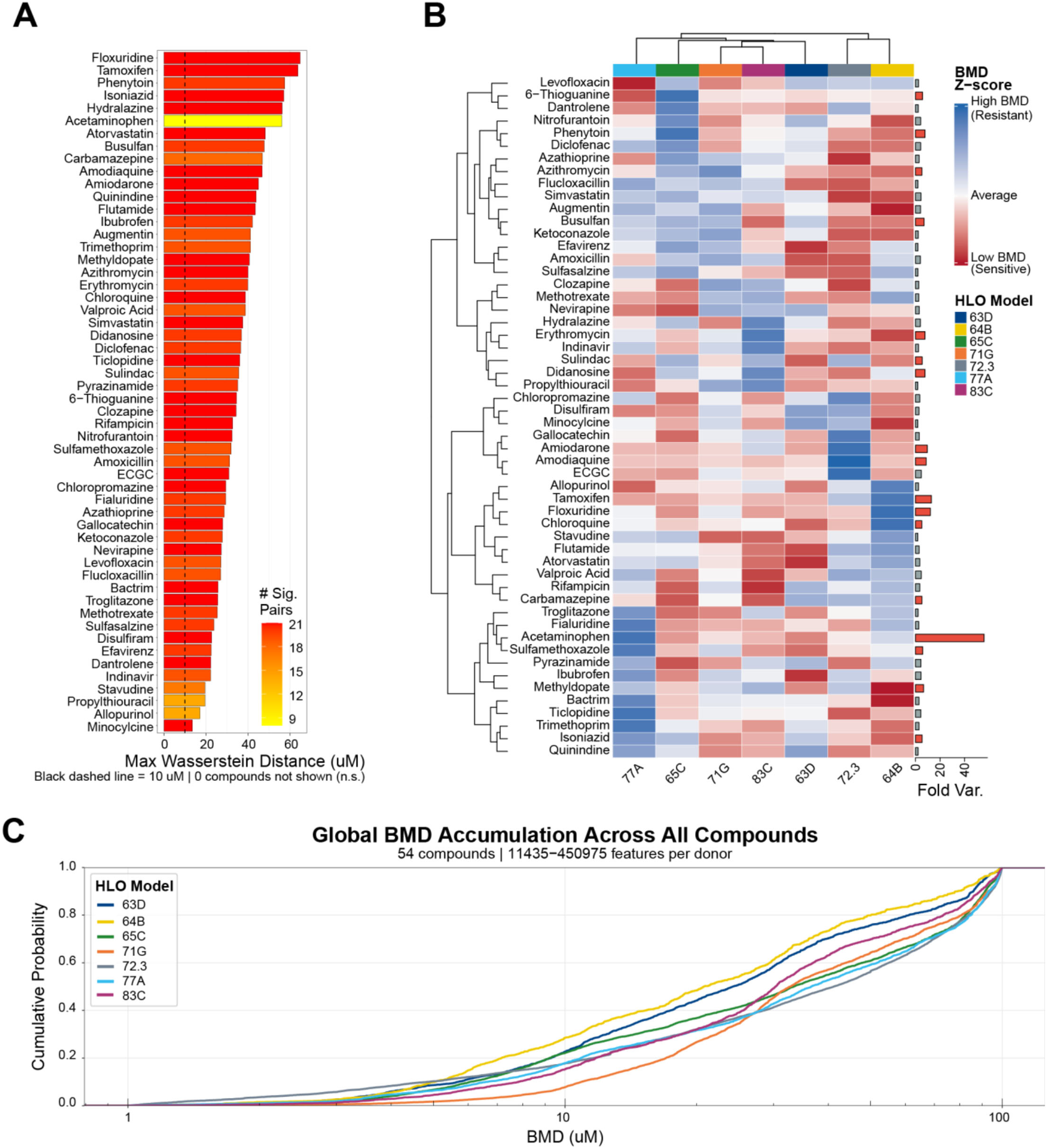
Compound-wide donor heterogeneity in benchmark dose responses across positive-control compounds. **(A)** Fifty-four DILI compounds with at least one significant donor difference, ranked by the maximum pairwise Wasserstein distance between donor-specific BMD distributions. Bar color denotes the number of significant donor pairs per compound; the dashed line marks 10 μM. **(B)** Hierarchical clustering heatmap of z-scored median BMD values across the same 54 compounds and seven donors. Red indicates below-average BMD (greater sensitivity) and blue indicates above-average BMD (greater resistance) relative to the compound-specific mean. Right-side bars show fold variability per compound (maximum/minimum donor median BMD), with red bars indicating >5-fold variability. **(C)** Global empirical cumulative distribution functions (eCDFs) of feature-level BMD values pooled across the 54 compounds for each donor (11,435–450,975 features per donor). All BMD, BMDL, and BMDU values were restricted to the tested dose range (≤100 μM) prior to analysis. Pairwise differences between donor distributions were assessed by Kolmogorov– Smirnov test with FDR correction.

classification analysis, exhibited substantial inter-donor variability in BMD-derived phenotypic responses, with feature-level BMD values spanning more than an order of magnitude across donors. This variability is consistent with the idiosyncratic nature of sulindac-associated hepatotoxicity, which has been linked to inflammatory co-stimulation pathways.^19–21^ Taken together, these results show that inter-individual variability in morphological dose-response is a pervasive property of hepatotoxic compound exposure, spanning conserved responses with overlapping donor distributions to highly divergent responses with non-overlapping BMD confidence bounds across the donor panel.

### Organoid Points of Departure Relative to Clinical Exposure

To ask whether these points of departure fall within the range of human exposure, we translated each compound’s BMC_50_ into a human-equivalent plasma concentration using an *in vitro* mass-balance model with the measured plasma free fraction and compared the result with curated literature C_max_ values for 68 of the 74 compounds, 66 of which had a curated free fraction (**Supplemental Figure S4; Supplemental Table S4**).

Modelling in-well drug adsorption lowered the free concentration by up to four orders of magnitude for lipophilic compounds such as amiodarone, tamoxifen and the statins, and left a residual offset of 23-fold (95% CI 12–45) between the organoid and clinical exposure distributions. After correction, the translated point of departure fell within 10-fold of clinical C_max_ for 41 of 66 compounds and within 100-fold for 60 of 66 (**Supplemental Figure S4B**). This correspondence was distributional rather than compound-specific: the weak positive rank correlation between translated point of departure and C_max_ (ρ = 0.30) was reproduced when BMC_50_ values were permuted with the physicochemical correction retained (ρ = 0.29) and disappeared once that correction was conditioned upon (partial ρ = 0.07), whereas simulation of an assay with genuine compound-specific translation predicted ρ = 0.87 under the same concentration range, censoring and inter-donor variability (**Supplemental Figure S4C**). Hepatocellular injury potency measured in this system is therefore statistically independent of clinical exposure, and the two must be combined explicitly rather than inferred from one another.

### Unsupervised Phenotypic Clustering Identifies Compound Groups Enriched for Shared Mechanistic Annotations

To investigate whether HLO morphological profiles encode not only DILI risk but also the underlying mechanism of hepatotoxicity, we performed channel-resolved unsupervised analysis on line 64B, selected as the most drug-sensitive HLO model based on BMD analysis (**Figure 6C**). Dose-level features were aggregated into a single vector of median values across five concentrations, generating a dose-response morphological profile for each compound. Leiden community detection on UMAP embeddings of the 54 DILI compounds identified four clusters with distinct phenotypic signatures (**Figure 7A**). Each cluster showed enrichment for features associated with specific imaging channels as evaluated by Shapley additive explanatory analysis (SHAP) (**Supplemental Figure S3**), which correspond to known hepatotoxicity mechanisms when cross-referenced with published pharmacology (**Figure 7B**; **Supplemental Table S2**).

**Figure 7.**
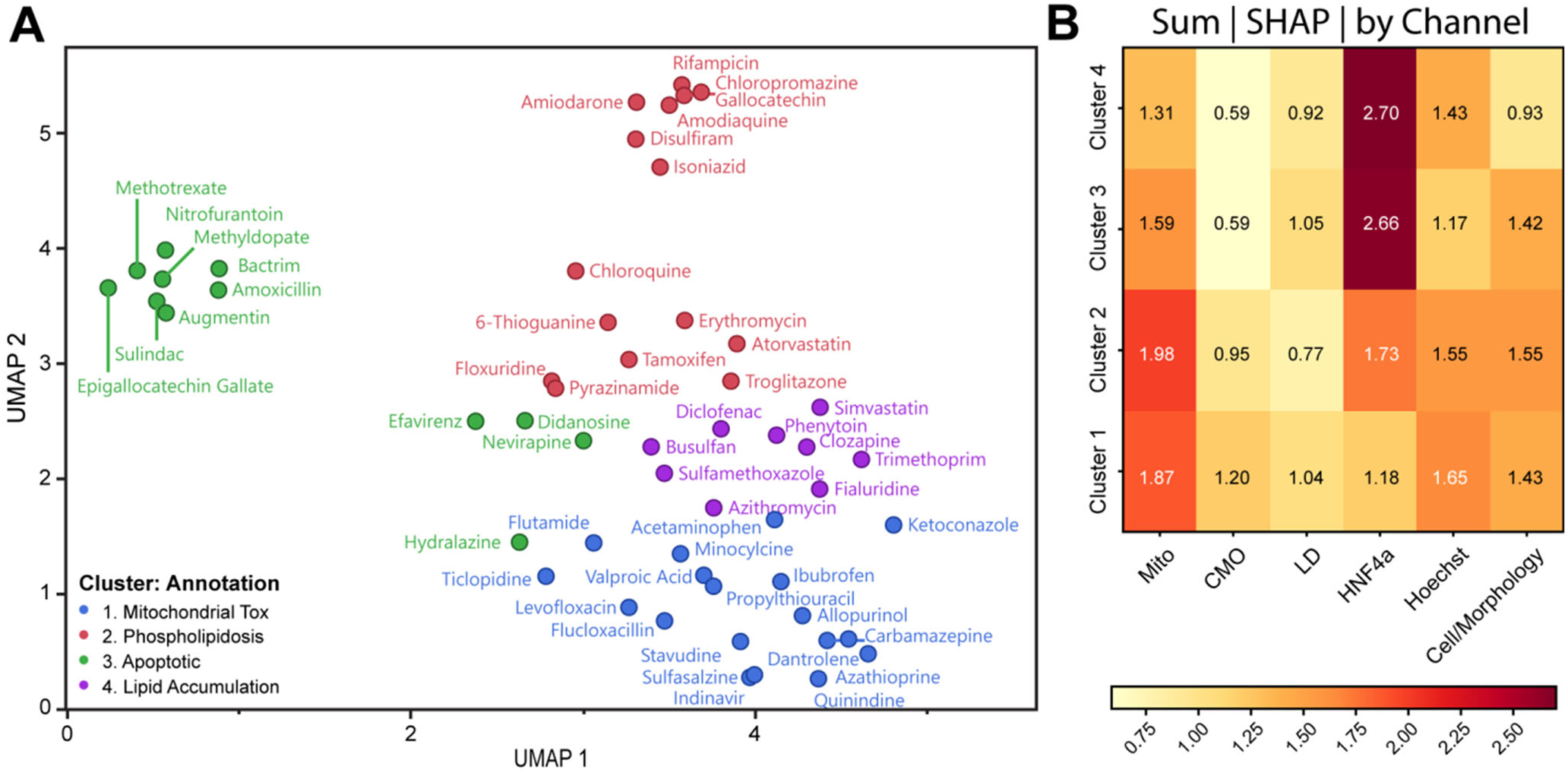
Phenotypic clustering reveals mechanism-associated hepatotoxicity signatures in HLOs. **(A)** UMAP projection of compound-level morphological profiles from line 64B HLOs, colored by cluster. Four clusters were identified and labelled by their enriched mechanistic annotation: mitochondrial perturbation (Cluster 1), cationic amphiphilic drug enrichment with membrane and lipid features (Cluster 2), nuclear and chromatin stress (Cluster 3), and lipid accumulation (Cluster 4). Cluster labels denote enrichment for compounds sharing published mechanistic annotations; the Cluster 2 phospholipidosis assignment is confirmed directly by phospholipid-specific staining (Supplemental Figure S5). **(B)** Channel-level contributions to cluster identity, quantified as the sum of absolute SHAP values per imaging channel. Mito, MitoTracker; CMO, HCS CellMask Orange; LD, lipid droplet; HNF4α, hepatocyte nuclear factor 4α; Hoechst, DNA stain; Cell/Morphology, cell-shape and texture features.

Cluster 1 (n = 18) was enriched for mitochondrial-associated features, indicating mitochondrial perturbation rather than demonstrated primary mitochondrial toxicity, and included known mitochondrial toxicants such as valproic acid, stavudine, flutamide, and ketoconazole, as well as acetaminophen. The concurrent presence of lipid-associated features is consistent with known links between mitochondrial dysfunction and secondary lipid accumulation.^22,23^ Cluster 2 (n = 15) exhibited strong multichannel perturbation, with CellMask (cytoplasm/membrane) features as the dominant signal, followed by lipid and HNF4α-associated features. This cluster was enriched for cationic amphiphilic drugs (CADs), including amiodarone, chlorpromazine, chloroquine, and tamoxifen, consistent with phospholipidosis, characterized by lysosomal phospholipid accumulation and altered membrane dynamics.^24–26^ Using a phospholipid-specific stain, amiodarone, chlorpromazine and tamoxifen each produced dose-dependent phospholipid accumulation at sub-cytotoxic concentrations (2.34-fold over vehicle at 89% of control nuclei, 5.04-fold at 87%, and 3.27-fold at 80%, respectively), whereas buspirone, a lysosomotropic cation that does not induce phospholipidosis, remained indistinguishable from vehicle across the full dose range (**Supplemental Figure S5, Supplemental Methods**). Three of the four cationic amphiphilic drugs in this cluster exhibit the phospholipidosis phenotype, confirming the phospholipidosis cluster annotation. Cluster 3 (n = 12) was defined by enrichment of Hoechst (nuclear/DNA) features and included compounds associated with immune-mediated or genotoxic stress, including hypersensitivity hepatitis, such as Augmentin (HLA-DRB1*15:01), nevirapine (HLA-DRB1*01:01), and the three most common causes of drug-induced autoimmune hepatitis: hydralazine, methyldopate, and nitrofurantoin.^27^ The nuclear-dominant signature is consistent with morphological features associated with chromatin condensation and DNA fragmentation (Supplemental Figure S3). Because the HLO system contains no adaptive immune cells, this nuclear signature cannot itself constitute evidence of an immune-mediated mechanism, and the co-clustering of hypersensitivity-associated drugs remains an unexplained association rather than a mechanistic assignment. Cluster 4 (n = 9) showed the lowest overall perturbation magnitude, with lipid droplet-associated features as the primary signal. This cluster included compounds linked to delayed-onset or metabolically mediated toxicity, such as fialuridine (a prototypical mitochondrial DNA inhibitor^28^), diclofenac (reactive metabolite-driven mitochondrial uncoupling^29^), and clozapine (metabolic idiosyncrasy^30,31^), consistent with more subtle early phenotypic changes in short-term in vitro assays.

These results indicate that HLO morphological profiles carry information beyond binary DILI classification, grouping compounds that share published mechanistic annotations. The observed correspondence between imaging features and known hepatotoxicity mechanisms provides a framework for generating hypotheses about the mechanism of action of novel compounds. Compounds profiled in this system can be contextualized relative to annotated reference clusters, enabling prioritization of mechanistic follow-up studies using targeted functional assays.

### TNFα Co-treatment Unmasks Sulindac-Induced Hepatotoxicity in the HLO Model

Sulindac is a clinically established hepatotoxicant whose DILI mechanism has been characterized as an immune-allergic hypersensitivity event or a metabolic idiosyncratic event that manifests after prolonged exposure, requiring pro-inflammatory co-stimulation to manifest injury in-vitro at clinically relevant exposures.^19–21^ Consistent with this mechanism, sulindac was misclassified as non-hepatotoxic in our standard screen, producing a dose-averaged ML DILI score of 0.30 ± 0.05, well below the classifier threshold of 0.38 (**Figure 3A**). Conventional viability assessment detected cytotoxicity only at 100 µM, a supraphysiologic concentration consistent with previous HepG2 reports (∼150 µM)^19^ and difficult to distinguish from general xenobiotic stress (**Figure 8A**). However, TNFα co-treatment (10 ng/mL) modestly potentiated sulindac-induced viability loss across the dose range. When HLOs were treated with sulindac sulfide, a sulindac metabolite which is the principal mediator of sulindac hepatotoxicity,^19,21^ pronounced dose-dependent cytotoxicity was observed, with TNFα co-treatment further enhancing injury and reducing viability to approximately 20% at 100 µM (**Figure 8B**).

**Figure 8.**
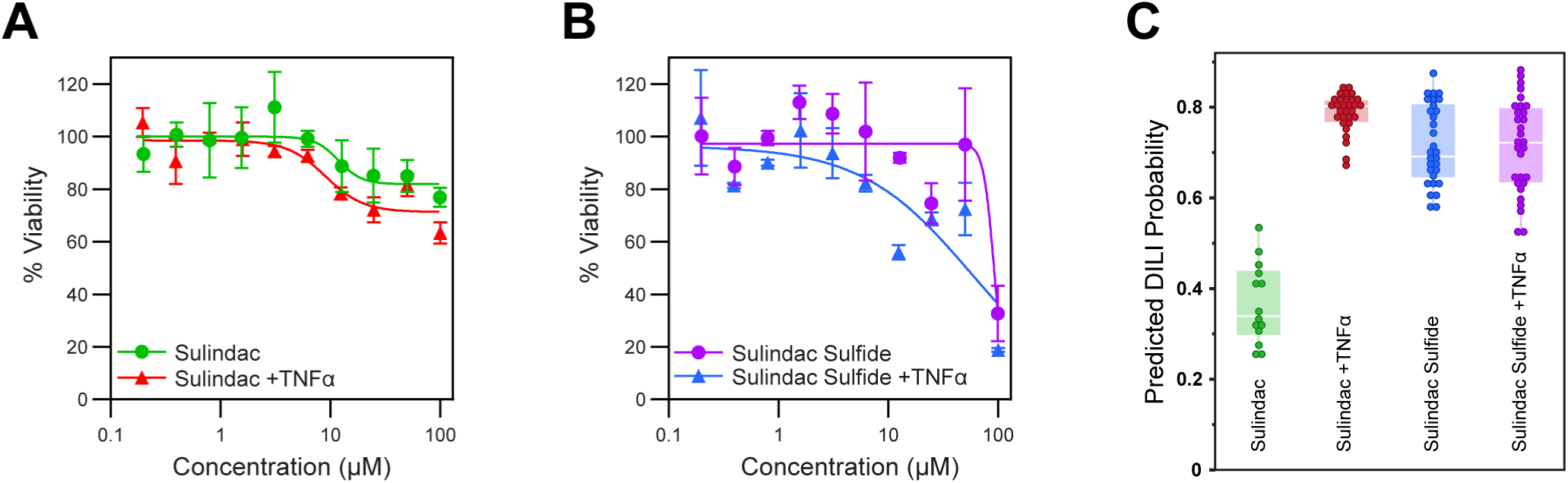
TNFα increases model-predicted DILI probability of sulindac in HLOs. **(A)** Dose–response viability curves for 72.3 HLOs treated with sulindac alone or in combination with TNFα (10 ng/mL). **(B)** As in (A), for the active metabolite sulindac sulfide ± TNFα. Data in (A) and (B) are mean ± SD. **(C)** Predicted DILI probabilities from the phenotypic machine-learning model for sulindac and sulindac sulfide with or without TNFα co-treatment. Each point represents one well; boxes denote the interquartile range, and the horizontal line indicates the median.

In an independent follow-up study, we assessed ML DILI probabilities in 72.3 HLOs treated with sulindac ± TNFα or sulindac sulfide ± TNFα across the same dose range and evaluated them using the previously trained DILI risk model for 72.3 HLOs. At the ML classification level, sulindac produced a DILI risk probability of 0.37, comparable to the previous screen, and both sulindac and sulindac sulfide produced significantly elevated DILI probability scores under TNFα co-treatment, with dose-averaged scores ranging from 0.71–0.78, effectively reclassifying sulindac from a false negative to a true positive (**Figure 8C**). Sulindac sulfide elicited elevated DILI scores even without TNFα, consistent with its identification as the direct hepatotoxic species. These results demonstrate that the HLO platform can detect inflammation-potentiated DILI mechanisms when supplemented with minimal pro-inflammatory priming. This approach enables identification of compounds with immune-mediated idiosyncratic hepatotoxicity that would otherwise escape detection under basal screening conditions.

## Discussion

The increasing emphasis on NAMs in drug safety assessment reflects a broader effort to improve the human relevance and translational fidelity of preclinical toxicology while accelerating drug development. Human organoid systems have emerged as promising platforms for modeling organ-specific toxicity; however, concerns regarding reproducibility, donor variability, and their ability to capture diverse mechanisms of DILI have limited widespread adoption. In this study, we demonstrate that patient-derived HLOs combined with high-content morphological profiling and supervised ML provide a robust and scalable platform for detecting phenotypic patterns associated with DILI with high sensitivity and specificity across genetically diverse donors. Despite substantial baseline phenotypic heterogeneity among seven HLOs, drug-induced responses were reproducible and interpretable. Importantly, consensus predictions across donors further improved predictive accuracy relative to any single line. These findings support multi-donor HLO phenotypic screening as a scalable NAM for early-stage hepatotoxicity hazard identification, enabling prioritization of scaffolds and compounds with reduced hepatocellular liability during hit-to-lead and lead optimization. However, prediction of human clinical DILI manifestation requires both accurate detection of DILI risk and accurate prediction of hepatic exposure, ideally within microphysiological systems incorporating flow to maintain drug concentration gradients. Consistent with this, several studies have reported increased DILI risk when serum drug concentration exceed ∼1.1 –1.3 µM.^32,33^

Our ML approach yielded compound-level AUROC values ranging from 0.88–0.91 across DILI patient-derived HLOs and achieved a consensus balanced accuracy of 89.7% (sensitivity 94.4%, specificity 85.0%). ML performance substantially exceeded conventional cytotoxicity readouts and matched or surpassed other in vitro DILI prediction platforms.^16,34,35^ ML enabled approximately a three-fold improvement in hepatotoxic compound detection compared to viability-based classification, reflecting its ability to capture early or sublethal phenotypic perturbations, including mitochondrial stress, lipid accumulation, and cell and nuclear morphological changes, that precede overt cell death. These findings underscore the need for phenotypic endpoints beyond simple viability measurements, as hepatotoxic injury often manifests initially as adaptive stress responses rather than acute cytotoxic collapse.

Several methodological advances contributed to the robustness of this predictive framework. Features were aggregated at the well level prior to modeling, reducing noise associated with single-cell variability while preserving compound-induced phenotypic signals. Compound-grouped cross-validation prevented information leakage between training and testing sets, and inverse-frequency weighting addressed class imbalance without synthetic oversampling. Plate-wise MAD robustization mitigated systematic plate effects, while automated ensemble model selection using AutoGluon enabled robust model optimization without manual hyperparameter tuning. Collectively, these refinements improved generalization performance and established a principled analytical framework for high-content toxicity screening.

A major strength of this study is the inclusion of HLOs derived from seven iPSC lines with six established from donors enrolled in DILIN, and one healthy donor (72.3). Baseline phenotypic characterization confirmed substantial donor-specific variation in hepatocyte morphology, lipid metabolism, and mitochondrial features; however, this heterogeneity did not impair predictive performance. Notably, the healthy donor line (72.3) achieved compound-level classification performance comparable to the best-performing DILI patient-derived lines (AUROC 0.89), demonstrating that DILI-associated phenotypic signatures can be captured even in non-diseased hepatocyte backgrounds. Pairwise prediction correlations revealed both highly concordant donor pairs (63D–65C, r = 0.85) and more divergent combinations (72.3–77A, r = 0.66), indicating that some lines contribute overlapping predictive information, whereas others provide complementary coverage of compound responses. These observations highlight an important conceptual advantage of multi-donor organoid systems: inter-line variability may capture distinct mechanistic pathways to hepatotoxicity. Consequently, consensus predictions across donors improve classification performance and are a step towards estimating population-level heterogeneity observed in clinical DILI. Thus, an efficient, maximally predictive, HLO-based hepatotoxicity screening platform may be constructed from multiple orthogonally responding lines to better approximate population-level DILI risk.

To contextualize these findings, we compared performance in HLOs to immortalized cancer cell lines. Although the same ML workflow improved classification performance relative to viability assays in HepG2, HUH7, HCT116, and U2OS cells, predictive accuracy remained substantially lower than that achieved in HLOs (80.5% vs. 89.4% for hepatotoxic compounds, respectively). The deficit was most pronounced for non-hepatotoxic compounds, where cancer cell lines exhibited higher false-positive rates. These differences likely reflect the altered baseline morphology, dysregulated metabolism, and limited xenobiotic biotransformation capacity characteristic of immortalized cells.^36^ In contrast, HLOs improved hepatocyte-like metabolic competence, enabling detection of physiologically relevant morphological responses to xenobiotic exposure.

Despite strong overall performance, several systematic misclassifications highlight important limitations of our HLO platform. Three persistent false negatives, Sulindac, Bactrim, and Augmentin, are associated with immune-mediated or idiosyncratic hepatotoxicity that requires non-parenchymal or adaptive immune interactions absent in the current HLO system. For example, Bactrim and Augmentin frequently induce immune-mediated DILI, while sulindac is associated with hypersensitivity-driven hepatotoxicity.^37^

Two anticipated limitations of the HLO model are capturing idiosyncratic immune-mediated DILI and detecting toxicity for compounds whose hepatotoxicity emerges only under chronic dosing or in combination therapies. To investigate these limitations, we examined inflammation-dependent hepatotoxicity using TNFα co-treatment. Sulindac did not elicit a significant response under basal conditions but produced pronounced phenotypic and viability changes when combined with TNFα stimulation. These findings demonstrate that the HLO platform can detect inflammation-potentiated hepatotoxicity under appropriate pro-inflammatory conditions. Incorporating cytokine-based perturbations into screening paradigms may therefore improve detection of inflammation-dependent mechanisms of idiosyncratic DILI and help distinguish primary hepatocellular injury from downstream immune-mediated effects.

The three persistent false positives (propranolol, pilocarpine, and miglustat) were consistently predicted as DILI-positive across all seven patient HLOs. This uniformity indicates that these compounds induce genuine hepatocyte phenotypic perturbations in the HLO system at the concentrations tested, even though they are not associated with clinical DILI under conventional dosing. All three were flagged well above their reported total C_max_ by approximately 66-fold for propranolol (therapeutic C_max_ ∼945 +/- 424 nM),^42^ 210-fold for pilocarpine and 23-fold for miglustat. These margins narrow considerably once exposure is anchored by applying the in-well dosimetry correction and the 23-fold static-to-flow offset described above leaves propranolol and pilocarpine approximately an order of magnitude above clinical exposure (margins of ∼12 and ∼9, respectively), but places miglustat at essentially its clinical Cmax (margin ∼1). Propranolol is ∼90% plasma protein bound and therefore gains substantially from the correction, whereas pilocarpine and miglustat are essentially unbound, so their nominal margins are consumed almost entirely by the static-culture offset. Propranolol and pilocarpine are accordingly better described as hepatocellular hazards detected outside the therapeutic window than as assay artifacts. The platform detects miglustat as a reproducible phenotypic perturbation at clinically relevant concentrations, and this likely represents a specificity limitation rather than one that exposure anchoring can resolve.

Compounds like diclofenac and nevirapine, which are associated with immune-allergic forms of DILI,^43,44^ were correctly identified as hepatotoxic in our assay. This suggests that some immune-mediated DILI may originate from primary epithelial or mesenchymal perturbations that are detectable in our HLO system. These findings demonstrate that the HLO platform can capture inflammation-potentiated hepatotoxicity when an appropriate inflammatory context is present. Incorporating cytokine-based perturbations into screening paradigms may therefore improve detection of inflammation-dependent mechanisms of idiosyncratic DILI.

Beyond binary classification, benchmark concentration modeling provided a quantitative framework for evaluating inter-line variability in phenotypic sensitivity. BMDExpress3 analysis revealed substantial donor-to-donor variability in morphological benchmark concentrations across the compound panel, with some compounds showing conserved responses and others exhibiting more than an order-of-magnitude difference in phenotypic sensitivity. These differences are consistent with the inter-individual variability observed clinically, which is driven by genetic, metabolic, and immunologic factors. The present design cannot, however, attribute them to donor genotype: each line was differentiated independently, so donor and differentiation batch are confounded, and the data therefore establish variability between donor-derived lines rather than patient-specific susceptibility. High-content morphological profiling detects early cellular perturbations and stress responses following xenobiotic exposure, representing phenotypic points of departure at which cellular homeostasis measurably deviates from baseline. Such perturbations do not necessarily progress to irreversible toxicity or clinical liver injury, as hepatocytes frequently mount adaptive responses, including metabolic reprogramming, mitochondrial remodeling, and altered lipid handling, within physiological exposure ranges. Accordingly, these morphological points of departure should be interpreted as indicators of hazard potential rather than direct predictors of clinical toxicity. In early drug discovery, this distinction is advantageous, enabling compound ranking, hit-to-lead prioritization, and lead optimization based on relative hepatotoxic liability. Feature-level benchmark concentrations provide quantitative phenotypic points of departure that can be integrated with pharmacokinetic exposure metrics, such as the C_max_/AUC, to estimate safety margins and population-level risk distributions. Translation to clinical DILI risk prediction will likely require integration with quantitative frameworks such as in vitro-to-in vivo extrapolation and physiologically based pharmacokinetic modeling, ideally in MPS systems. These morphological benchmark concentrations provide quantitative phenotypic points of departure that capture inter-line differences in cellular stress sensitivity and, when integrated with pharmacokinetic exposure and IVIVE/PBPK modeling, may support estimation of hepatotoxic safety margins without implying direct prediction of clinical DILI.

Importantly, we observed and corrected the general reduction in apparent potency in 384-well static culture, which the present calibration allows us to partition into two components: loss of freely dissolved compound to the culture system, and collapse of the concentration gradient driving delivery into the organoid in the absence of replenishment. First, with high surface-area-to-volume ratio and 50 ‡L of medium in 384-well plates, a substantial fraction of compound partitions into plate plastic and cellular lipid rather than remaining freely dissolved. Modelling this explicitly reduced the estimated free concentration relative to nominal by up to four orders of magnitude for the most lipophilic compounds, and accounting for plastic and cellular binding beyond medium albumin alone lowered the panel-wide offset by a further 2.2-fold. Second, drug exposure in static culture systems reduces effective concentrations due to drug metabolism and collapse of the concentration gradient that drives delivery into the organoids. This analysis quantified a 23-fold (95% CI 12-45) reduction in potency in the assay, allowing estimation of whether the identified risk is in the range of human exposure.

In addition to dose-response modeling, unsupervised analysis uncovered distinct clusters of phenotypic perturbations that aligned with annotated DILI mechanisms.^45,46^ While clustering was broadly preserved across donors, modest differences in compound groupings reflect donor-specific sensitivities. These phenotypic signatures provide a foundation for mechanism-driven hypothesis generation and may support development of biomarker panels for stratifying hepatotoxic liability or identifying line-specific differences in sensitivity. Targeted follow-up using phospholipid-specific staining showed dose-dependent phospholipid accumulation at sub-cytotoxic concentrations for the cationic amphiphilic drugs of the phospholipidosis cluster, while a lysosomotropic non-inducer remained at vehicle levels (**Supplemental Figure S5**).

Finally, this work highlights several opportunities for further development of the platform. Extension to microphysiological systems incorporating multicellular architecture and physiologically relevant exposure dynamics could improve modeling of chronic or adaptive responses. Integration of immune components may enhance detection of idiosyncratic DILI, and incorporation of pharmacogenomic annotation could support personalized risk assessment.

In conclusion, multi-donor human liver organoids combined with high-content morphological profiling and ML provide an improved platform for detecting hepatotoxicity-associated phenotypes while capturing inter-line variability in drug responses. The system enables sensitive detection of early phenotypic perturbations, supports mechanistic hypothesis generation, and allows donor-line-specific response profiling in a human-relevant context. Continued integration with advanced in vitro models and quantitative translational frameworks will further enhance predictive performance and support safer drug development while reducing reliance on animal testing.

## Limitations of this Study

This study has several limitations. First, the compound library of 74 drugs, although selected for high-quality clinical DILI annotation, is small relative to the chemical space relevant to drug discovery, and model performance may degrade on larger or more structurally diverse panels. The Youden-optimal consensus threshold was tuned on the same 74-compound dataset used for evaluation; independent validation on a held-out compound set will be required to establish generalizability. Second, although the HLO platform contains hepatic stellate and Kupffer cells,^10^ it lacks immune cell populations, limiting detection of HLA-restricted hypersensitivity and other idiosyncratic, immune-mediated DILI mechanisms. The persistent false negatives associated with immune-mediated injury (sulindac, Bactrim, Augmentin) illustrate this constraint. TNFα co-treatment with sulindac recovers detection by simulating an inflammatory milieu, but it is not a substitute for an antigen-specific adaptive immune response. Third, the screen used acute (72 h) exposure conditions that can miss chronic toxicity phenotypes, such as those driving methotrexate hepatotoxicity. Translation of drug exposure in static culture systems to clinical pharmacokinetics requires integration with quantitative in vitro-to-in vivo extrapolation (QIVIVE) and physiologically based pharmacokinetic modeling, which may be more effectively implemented in microphysiologic systems (MPS) with active flow that better approximate in vivo exposure kinetics. Fourth, each donor line was differentiated independently, so donor genotype and differentiation batch are confounded in this dataset. Independent differentiation replicates were not performed and a single non-DILI control line cannot support a case-control inference, so the inter-line differences reported here establish variability between donor-derived models rather than patient-specific susceptibility. Attributing that variance to donor genotype will require systematic repeat differentiations across lines and a larger control cohort. Fifth, the binary DILI labels used for training collapse compounds that differ substantially in incidence, severity, latency, mechanism, and intrinsic versus idiosyncratic etiology. Six of the twenty negative controls are classified as Ambiguous-DILI-concern in DILIrank 2.0, so reported performance is sensitive to label stringency and should be interpreted alongside the annotation provided in **Supplemental Table S2**.

## Experimental Procedures

### Human Induced Pluripotent Stem Cell (iPSC) Generation

iPSC lines 63D, 64B, 65C, 71G, 77A, and 83C were derived from patient peripheral blood mononuclear cells in collaboration with the University of Michigan Human Stem Cell and Gene Editing Core, as previously described.^47^ Briefly, erythroid progenitors were isolated, expanded, and reprogrammed using episomal plasmids with a commercial erythroid progenitor reprogramming kit (StemCell Technologies, 05924). All patient enrollment, iPSC generation, and human liver organoid (HLO) differentiation were conducted under the approved institutional study protocol, HUM00204307, and all stem cell work was performed with the approval of the University of Michigan Human Pluripotent Stem Cell Research Oversight Committee (HPSCRO Record #1134). A control iPSC line (72.3) was obtained from Cincinnati Children’s Hospital from a healthy donor with no history of liver disease using the same protocol.

### Genomic Integrity Assessment and Cell Line Authentication

Genomic integrity and identity of each iPSC line were verified using SNP DNA microarray analysis. Genomic DNA from early-passage cells was isolated using the DNeasy kit (Qiagen) and processed by the University of Michigan Advanced Genomics Core using the Infinium Global Screening Array-24 v3.0 platform. Array data were analyzed using GenomeStudio CNV Analysis (CNV Partition v3.2.0); copy-number variants were filtered by confidence score, loss of heterozygosity, and manual inspection of B-allele frequency and Log R ratio plots. All cell lines were identified as unique, and one confirmed CNV was identified: clone 71G contained an approximately 0.5-Mbp duplication (chr7:33,077,502–33,579,232) spanning most of the BBS9 gene.

### Human Liver Organoid (HLO) Differentiation

iPSC lines were differentiated into HLOs using a previously described protocol.^10,11^ Briefly, iPSCs were seeded at 1.3 × 10⁶ cells per well on growth factor–reduced Matrigel–coated (Corning, 354230) 6-well plates (VWR, 10062-892). Cells were induced to definitive endoderm with BMP-4 and Activin A (R&D Systems, 314-BP, 338-AC) for 24 hours, followed by Activin A alone for 2 days, and FGF-4 (PeproTech 100-31) ^48^ for 3 days. Definitive endoderm aggregates were embedded in 50 µL Matrigel droplets (Corning, 354234) within 6-well plates and treated with retinoic acid for 4 days. Organoids were then cultured for 12 days in hepatocyte growth medium (Lonza, CC-3198) supplemented with 10 ng/mL HGF (PeproTech, 100-39), 20 ng/mL oncostatin M (PeproTech, 300-10), and 100 nM dexamethasone (Sigma, D4902).

### HLO Dissociation and Compound Treatment

HLOs were dissociated using mechanical pipetting in combination with TrypLE (Gibco, 12604013) incubation for 10 minutes. Cells were pelleted by centrifugation at 500 × g for 5 minutes and washed 3 times with DMEM/F-12 (Gibco, 11320033). Following resuspension, cells were seeded at 8000 cells/well into collagen I-coated, black-walled, clear-bottom 384-well plates (Revvity, 6057700) in 50 µL medium containing 10 µM Y-27632 (Cayman Chemical, 10005583). After 24 hours, wells were treated with compounds at 5 concentrations spanning 1-100 µL for 72 hours.

### Compound Library and Annotation

A library of 74 compounds was curated to include clinically relevant drugs with and without documented hepatotoxicity (**Supplemental Table S2**). Fifty-four compounds were designated as positive controls (PC) for DILI: 45 selected by LiverTox scores A–C, two catechin-class hepatotoxicants (epigallocatechin gallate and gallocatechin; Green Tea, LiverTox score A), and two non-US-FDA-approved hepatotoxicants (fialuridine and flucloxacillin; see **Supplemental Table S2** for individual rationales).^49–51^ Five additional compounds with well-characterized hepatotoxic mechanisms (chloroquine, clozapine, indinavir, tamoxifen, and troglitazone) served as mechanistic standards. Twenty compounds were designated as negative controls (NC), including drugs with low hepatotoxic potential (LiverTox score E), compounds with established clinical safety, or drugs absent from major DILI reference databases. DMSO served as the vehicle control. Classifications were cross-referenced against the DILIRank 2.0 and DILIst datasets from the U.S. FDA Liver Toxicity Knowledge Base, the LiverTox Master List of Drugs, and the HESI OASIS Consortium compound reference list.^15,52,53^ All compounds were sourced from commercial vendors and solubilized as 10 mM stock solutions in fresh DMSO in echo-compatible 384-well plates and were stored at room temperature in a desiccator. Compounds were transferred to intermediate dilution plates with a Beckman Echo 650 in triplicate, with DMSO concentrations normalized across wells and were randomized to minimize plate artifacts.

### TNFα Co-treatment Experiments

To investigate sulindac-induced hepatotoxicity under inflammatory conditions, HLOs were pre-treated with recombinant human TNFα (10 ng/mL; PeproTech, 300-01A) for 2 hours prior to compound addition, and TNFα was maintained throughout the 72-hour drug incubation period. Sulindac was tested using the same 5-point, 1:3 serial dilution scheme used in the standard screen. DMSO vehicle and non-hepatotoxic control compounds were treated with TNFα to account for baseline inflammatory effects. ML DILI scores were computed using the trained AutoGluon models and compared between TNFα co-treated and standard conditions using the previously described consensus prediction framework.

### Cell Fixation and Staining

Five fluorescent channels were collected in two imaging passes to label nuclei, cells, neutral lipid droplets, mitochondria, and HNF4α. After 72-hour drug exposure, cells were incubated with 250 nM MitoTracker Deep Red (Invitrogen, M22426) in HGM for 45 minutes, then fixed and permeabilized with 4% paraformaldehyde and 0.3% Triton X-100 for 15 minutes. Cells were blocked for 1 hour at room temperature (RT) in 5% Bovine Serum Albumin (BSA; Sigma Aldrich, A7030-100G) before being incubated with primary anti-human HNF4α antibody (Invitrogen, MA5-32295) in 1% BSA, 0.3% Triton X-100 overnight at 4°C, in the dark. Alexa Fluor 594 goat anti-rabbit secondary antibody (1:1,000; Invitrogen, A11012) was incubated in 1% BSA and 0.3% Triton X-100 along with Hoechst 33342 (1:1,000; Invitrogen, 62249) and HCS LipidTox Green (1:1,000; Invitrogen, H34475) for 1 hour, RT, dark. Cells were washed three times with PBS in between steps. After imaging, cells were overstained with HCS CellMask Orange (1:10,000; Invitrogen, H32713) for 45 minutes at RT, dark and plates were re-imaged. HCS CellMask Orange images were aligned to first-pass images using a custom Python pipeline by alignment of nuclear masks from each pass.

### High-Content Imaging and Analysis

Nine fields of view (FOVs) per well were acquired in 3D (10 µm depth, 2 µm intervals) using a 20X/1.0 NA water-immersion objective lens on the Yokogawa Cell Voyager 8000 high-content imaging system with a dual-microlens spinning disc confocal unit with a 50 µm pinhole. Maximum intensity projection images were produced to ensure optimal focus for every cell observation.^54^ FOVs were arranged in 3×3 tiles with 50-pixel overlap and stitched using Yokogawa CellPathfinder software (vR3.08.01). Images were captured on a 2k Hamamatsu Orca sCMOS camera (pixel size: 0.3125 µm), with exposure settings optimized per channel to maximize signal-to-noise ratio, avoiding saturation.

### Image Quantification and Analysis

Segmentation of regions of interest (ROI) was performed for nuclei, lipid droplets, and mitochondria, using Cellpose 3.0/cyto3. Raw images and segmentation masks were imported into CellProfiler 4.2.8.^55–57^ The cell and cytoplasm ROIs were identified in CellProfiler via propagation from the nucleus object. Features extracted included area/shape, intensity, intensity correlations, counts, and texture for all channels in all ROIs. Mitochondrial network morphology was quantified wherein segmented mitochondria were reduced to single-pixel-wide representations via the MorphologicalSkeleton module, followed by the AnalyzeSkeleton method to extract branching metrics including the number of branch points, endpoints, and mean branch length per cell. Lipid droplet and mitochondrial-derived features were summarized at the cell-level.

### Feature Processing and Normalization

High-content imaging generated single-cell morphological feature vectors for each treatment condition. Missing values arising from absent cellular compartments or segmentation artifacts were imputed using the per-plate median. Single-cell features were aggregated by computing the median across all cells within each well, generating one phenotypic profile per well; zero-variance features were removed. Well-level phenotypic profiles were normalized per plate using the median absolute deviation (MAD) robustization implemented in pycytominer^58^ using the DMSO vehicle controls, correcting for plate-to-plate technical variation while preserving treatment effects. The natural logarithm of treatment concentration (log-dose) was appended as an additional feature.

### Phenotypic ML Modeling

Phenotypic ML classifiers were trained independently for each of seven HLO donor lines using AutoGluon Tabular AutoML (v1.4) with the best_quality preset and AUROC as the optimization metric. To address class imbalance between hepatotoxic (54) and non-hepatotoxic (20) compounds, inverse-frequency sample weights (scikit-learn) were applied at the well level. DMSO vehicle control wells were excluded from model training. To prevent data leakage, compound-grouped five-fold cross-validation (GroupKFold) was used, ensuring all wells from a given compound were assigned to the same fold. With 1,110 treated wells per line and GroupKFold by compound (74 groups), each fold contains approximately 15 compounds (225 wells for training). OOF predicted probabilities of DILI were generated for each well exclusively by models trained on other compounds. Model performance was evaluated using AUROC, balanced accuracy, and F1 scores. Analysis notebooks/scripts are available on GitHub: https://github.com/SextonLab/iDILI-Predict.

### Cross-Donor Consensus Prediction

To derive per-drug consensus DILI predictions across donors, the arithmetic well-level OOF mean probability across the seven lines was computed as a global consensus metric. An optimal binary classification threshold was determined by maximizing Youden’s J statistic on the receiver operating characteristic (ROC) curve calculated using the compound-level OOF predictions and ground-truth DILI labels. Pairwise Pearson correlations of compound-level OOF probabilities were computed across all donor lines to quantify inter-line concordance. Hierarchical clustering (average linkage; distance = 1 − r) was applied to the resulting correlation matrix to identify groups of phenotypically similar donor lines.

### Benchmark Concentration (BMC) Modeling with BMDExpress3

Benchmark concentration (BMC) modeling was performed using BMDExpress3 (v3.20.14). Preprocessed MAD-robustized well-level phenotypic feature data were modeled separately for each Donor × Condition × Chemical combination with matched DMSO controls. Concentration-response relationships were fitted with Hill, Power, Linear, Polynomial (2°–4°), and Exponential (3, 5) models. The benchmark response was set to 1.021 standard deviations (5% relative to control) using the standard deviation benchmark type, with profile likelihood for BMDU/L estimation and a confidence level of 0.95. Best-fit models were selected by nested chi-square test followed by lowest Akaike information criterion (AIC). Hill models were flagged if the k parameter was less than 1/3 of the lowest positive dose; flagged models were replaced by the next best model with p-value > 0.05 (fraction of minimum BMD = 0.5). This yielded a single best-fit BMD value per feature per donor–condition– chemical combination.

### Exposure-Anchored Translation of Benchmark Concentrations

Compound-level BMC_50_ values were translated into human-equivalent plasma concentrations in three steps. First, the freely dissolved concentration in the well was computed with the Armitage *in vitro* distribution model as implemented in the httk R package (v2.7.4), parameterized for a 384-well plate with 50 µL working volume and 8,000 cells per well at pH 7.4, with medium albumin set to BSA at 20–50-fold below serum albumin (geometric midpoint 1.27 g/L) and serum lipid set to zero to reflect serum-free hepatocyte medium.

Physicochemical parameters (log K_ow_, aqueous solubility, melting point, Henry’s law constant, pK_a_) were taken from the httk chemical library for 61 of 74 compounds and computed with RDKit (Crippen log K_ow_; ESOL aqueous solubility) for the remaining 13. Second, under the free-drug hypothesis the human-equivalent total plasma concentration was computed as the product of BMC_50_ and the ratio of the free fraction in medium to the measured free fraction in plasma. Third, the offset between this quantity and clinical C_max_ was estimated as the geometric mean ratio across compounds, with 95% confidence intervals obtained by bootstrap (5,000 resamples). Whether per-compound pairing was tighter than chance was tested by permuting the C_max_ assignment 20,000 times; the contribution of the organoid measurement specifically was isolated by permuting BMC_50_ while retaining the physicochemical correction, and by partial correlation conditioned on that correction. Statistical power was assessed by simulating an assay with perfect per-compound translation subject to the observed concentration range, censoring and inter-donor variability. Plasma free fractions were curated from regulatory labels or primary pharmacokinetic literature; three compounds were assigned the value of a close structural analogue or drug class and results are reported both with and without them.

### Unsupervised Drug Clustering and Feature Prioritization

To identify patterns of phenotypic similarity among compounds across dose conditions, we performed unsupervised clustering on aggregated morphological profiles of normalized single-cell features. For each compound, median feature values were calculated at each concentration, and a pivoted feature matrix (long-to-wide transformation) was constructed with one row per compound and one column per concentration-feature combination. This transformation allowed us to capture dose-dependent phenotypic trends within each compound profile. Prior to dimensionality reduction and clustering, features were standardized using StandardScaler, and missing values were imputed using the median of each feature column. The resulting matrix was subjected to UMAP (umap-learn: 0.5.7) for dimensionality reduction (metric=’cosine’, n_neighbors=5, and min_dist=0.1) to preserve local structure among compound profiles. A k-nearest neighbor graph was constructed in UMAP space (n_neighbors=15) and Leiden clustering (leidenalg: 0.10.2) was performed at resolution 0.8 to detect communities of compounds with similar morphological effects.

## Abbreviations

APAP: Acetaminophen
AUC: Area under the curve
AUROC: Area under the receiver operating characteristic curve
AUPRC: Area under the precision recall curve
CYP: Cytochrome P450
DILI: Drug-induced liver injury
DILIN: Drug-induced liver injury network EGCG: Epigallocatechin gallate
HLO: Human liver organoid
iPSC: Induced pluripotent stem cell
MAD: Median Absolute Deviation
ML: Machine Learning
NAM: New approach methodologies
NC: Negative control
PC: Positive control
ROC: Receiver operating characteristic

## Acknowledgements

Research reported in this publication was supported by the National Institute of Diabetes and Digestive and Kidney Diseases (NIDDK) of the National Institutes of Health under award numbers U01DK065211 (Indiana University), U01DK065184 (University of Michigan), U01DK065201 (University of North Carolina-Chapel Hill), U01DK083020 (University of Southern California), U01DK083027 (Thomas Jefferson University/Albert Einstein Medical Center), and U24DK065176 (Duke University). Additional support is provided by the intramural programs of the NIDDK and National Cancer Institute. The content is solely the responsibility of the authors and does not necessarily represent the official views of the National Institutes of Health. We thank the Drug-Induced Liver Injury Network (DILIN) for their continued collaboration in study design and for enrolling patients for iPSC reprogramming. We thank the University of Michigan Human Stem Cell and Gene Editing Core Facility (RRID:SCR_026755) for assistance in generating iPSC lines, and the laboratory of Jason Spence for continued collaboration and assistance with iPSC differentiation. We are grateful to Patrick Maher for helpful advice on machine learning training. This work was supported by the National Institutes of Health (S10 OD034245 to J.Z.S.; R01 GM152417 to J.Z.S. and R.J.F.; and U01 DK065184 to R.J.F. and J.Z.S.). At the time of experimentation, S.R.M. was supported by T32 GM140223 and N.Z. was supported by T32 GM141746. All authors had access to all data and have reviewed and approved the final manuscript.

## Supplemental Figures

**Supplemental Figure S1.**
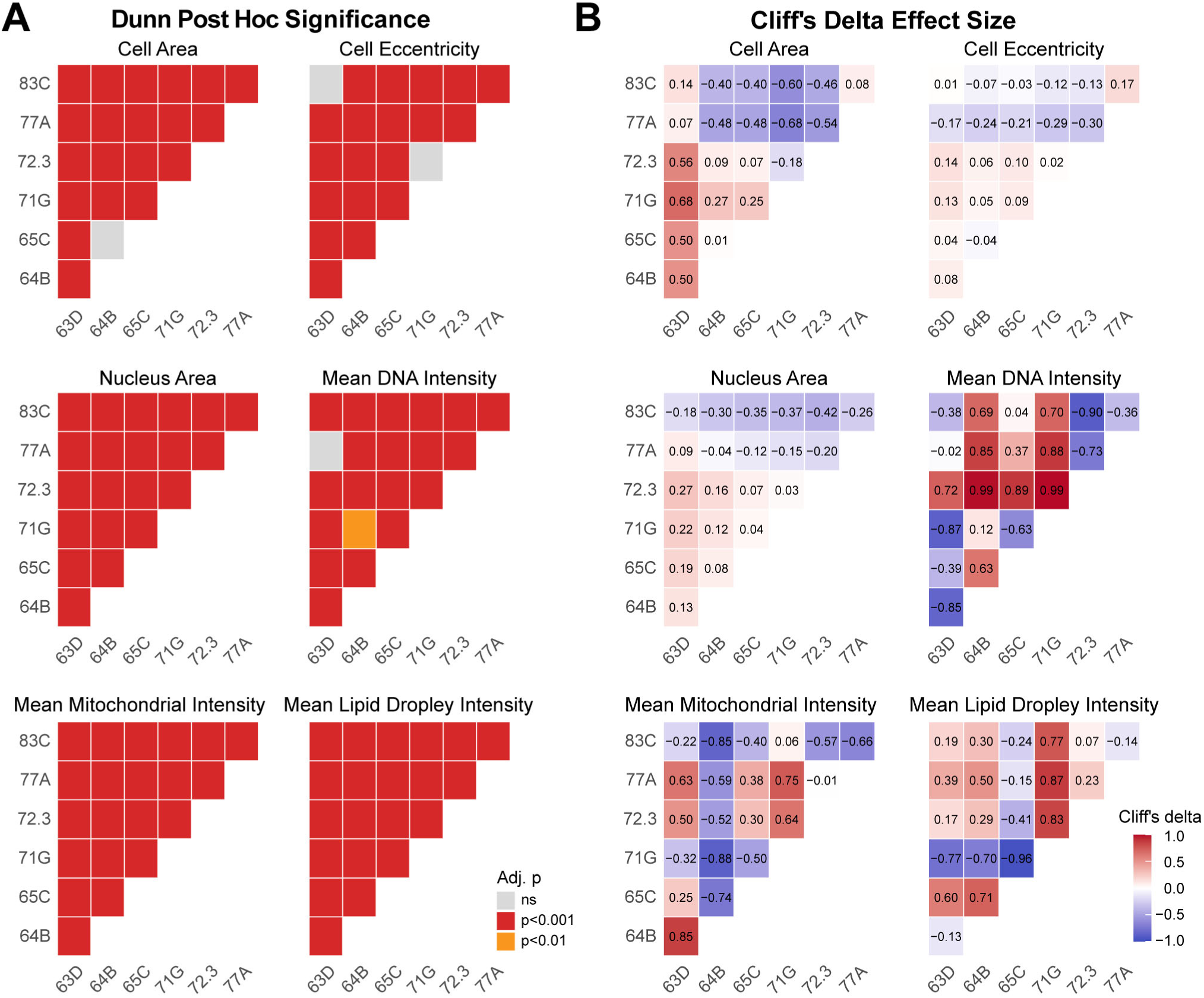
Pairwise statistical comparisons of baseline cellular features across HLO models. **(A)** Heatmaps showing pairwise differences among HLO models for representative cellular features. Statistical significance was assessed using Kruskal–Wallis testing followed by Dunn’s post hoc multiple-comparison analysis with Benjamini–Hochberg adjusted *p* values. Gray indicates non-significant differences, whereas warmer colors indicate increasing statistical significance. **(B)** Heatmaps of Cliff’s delta effect sizes for the same pairwise comparisons. Cliff’s delta quantifies the magnitude and direction of differences between groups. Red indicates higher values in group 1 relative to group 2, blue indicates lower values, and white indicates similar distributions. Values within tiles denote Cliff’s delta estimates, highlighting donor-dependent baseline phenotypic differences across HLO models.

**Supplemental Figure S2.**
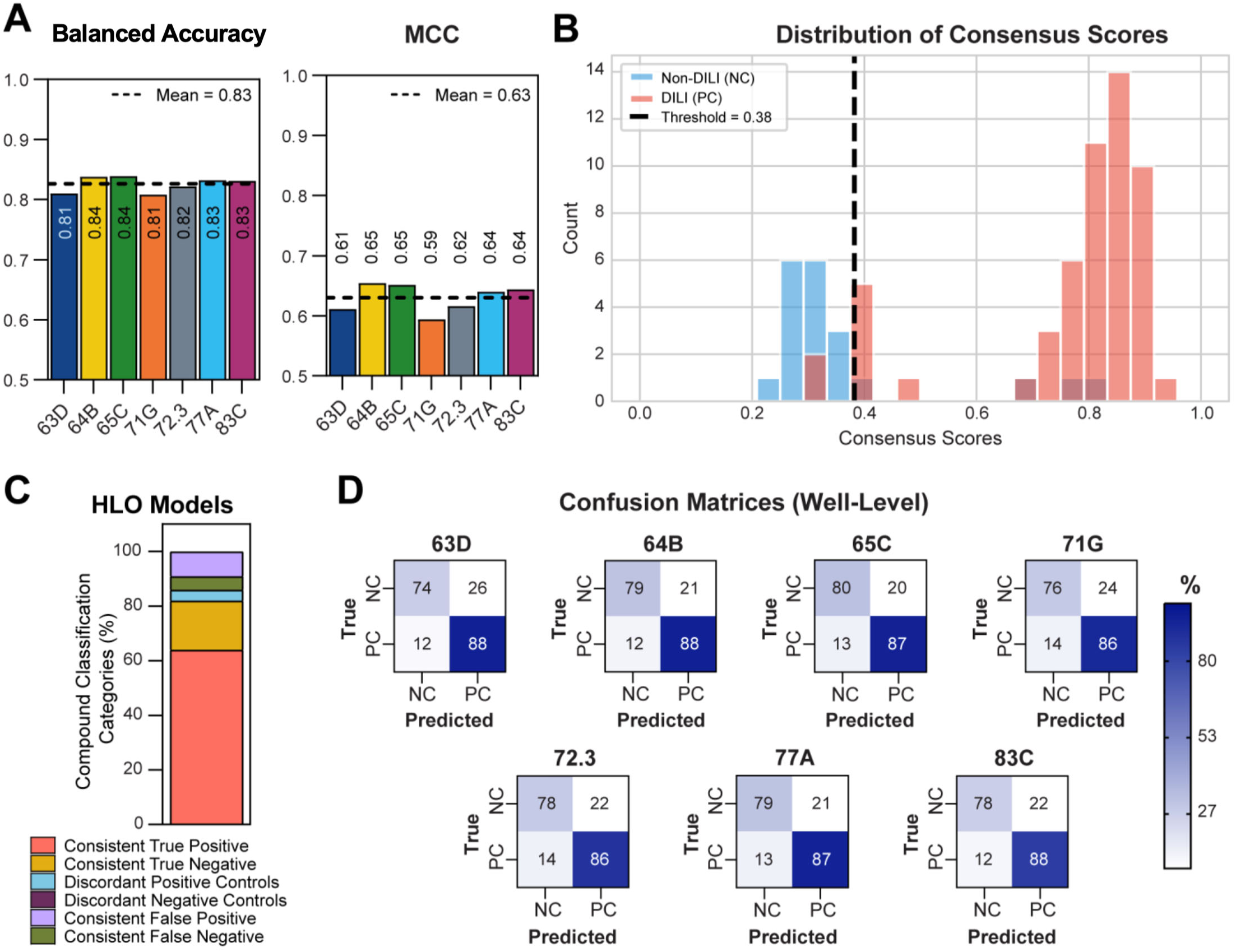
Well-level performance and classification metrics across individual and consensus HLO models. **A)** Balanced accuracy (left) and MCC (right) for each HLO model. These metrics demonstrate consistent predictive performance across genetically diverse donors. **(B)** Distribution of consensus DILI scores at the well level, showing separation between non-hepatotoxic (negative) and hepatotoxic (positive) wells. **(C)** Well-level compound classification categories for each HLO model and the consensus model, indicating the proportion of correctly classified hepatotoxic and non-hepatotoxic wells. **(D)** Confusion matrices calculated at the well level for all seven HLO models, highlighting sensitivity, specificity, and overall accuracy.

**Supplemental Figure S3.**
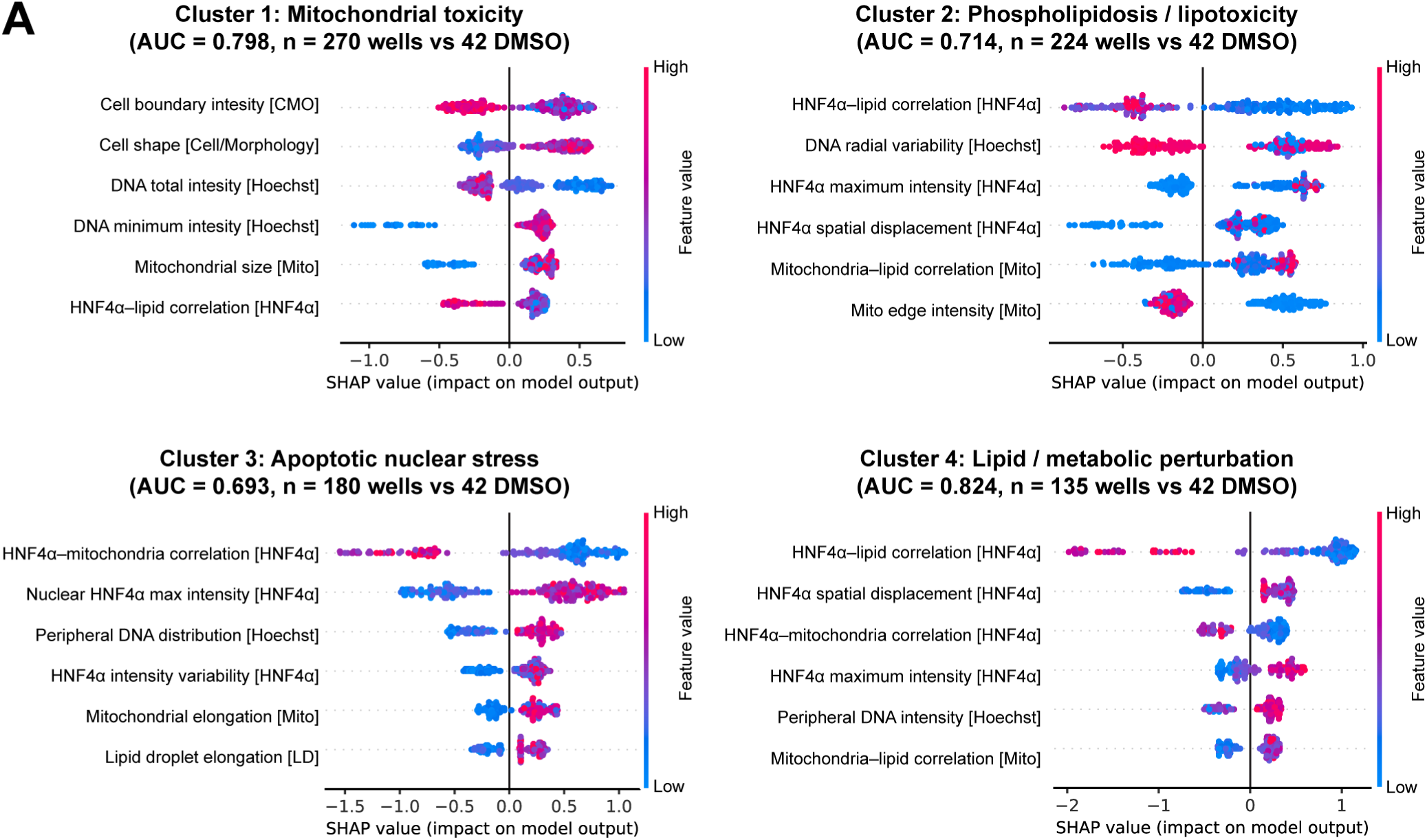
SHAP-based interpretation of mechanistic hepatotoxicity clusters identified from HLO morphological profiling. SHAP (SHapley Additive exPlanations) summary plots showing the top contributing morphological features driving classification of each phenotypic cluster identified from line 64B HLOs. Each point represents an individual well, colored by normalized feature value (blue = low, pink = high). Positive SHAP values indicate features contributing toward cluster assignment. Cluster 1 was characterized primarily by mitochondrial morphology and DNA intensity features, consistent with mitochondrial toxicity. Cluster 2 showed strong contributions from lipid-, membrane-, and HNF4α-associated features, consistent with phospholipidosis/lipotoxicity. Cluster 3 was enriched for nuclear organization and chromatin-associated features, consistent with apoptotic or nuclear stress phenotypes. Cluster 4 was defined by subtler HNF4α-, lipid-, and mitochondria-associated spatial relationship features, consistent with delayed-onset metabolic idiosyncratic toxicity. AUC values and the number of cluster-associated wells used for model training are shown for each classifier.

**Supplemental Figure S4.**
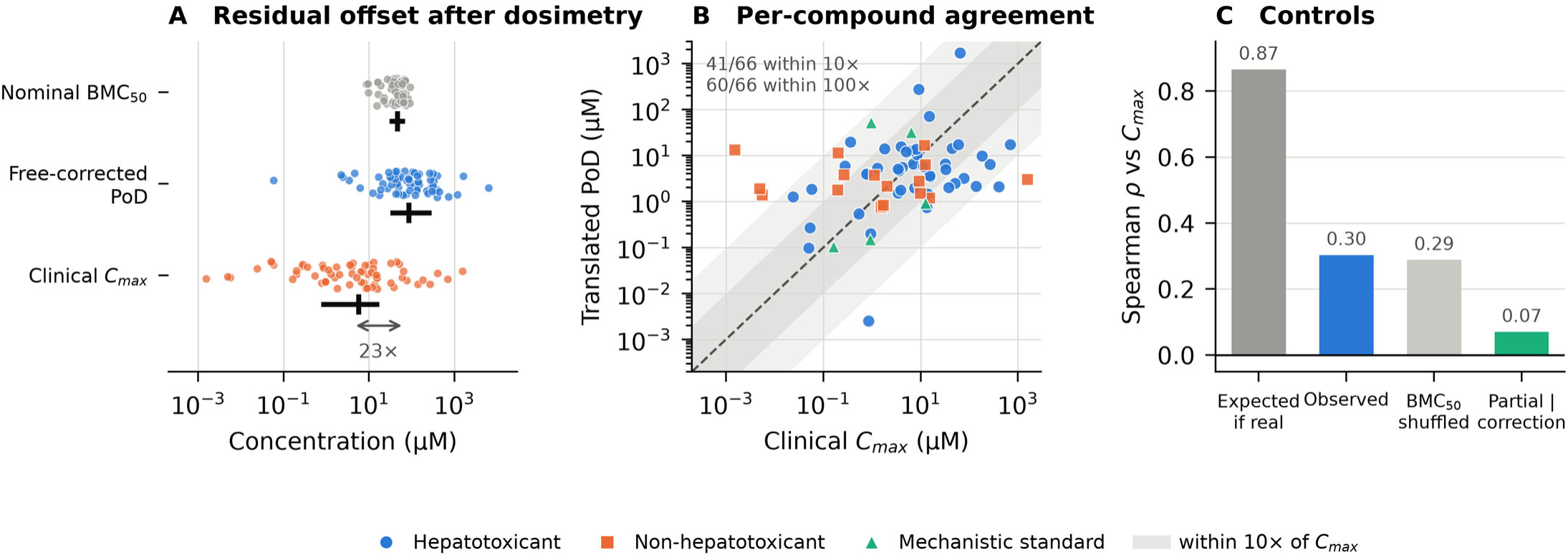
Organoid points of departure relative to clinical exposure. **(A)** Distributions of nominal BMC_50_, the free concentration after in-well dosimetry, and clinical total C_max_ for the 66 compounds with both a curated C_max_ and plasma free fraction. Points are individual compounds; bars mark median and interquartile range. The residual offset between the free-corrected point of departure and clinical exposure is 23-fold (95% CI 12–45). **(B)** Translated point of departure against clinical C_max_ for each compound. Dashed line, identity; shaded bands, within 10-fold and 100-fold of C_max_. **(C)** Control analysis. The observed rank correlation is reproduced when BMC_50_ is permuted with the physicochemical correction retained, and vanishes when conditioned on that correction, whereas an assay with genuine compound-specific translation would be expected to give ρ = 0.87 under the same concentration range, censoring and inter-donor variability.

**Supplemental Figure S5.**
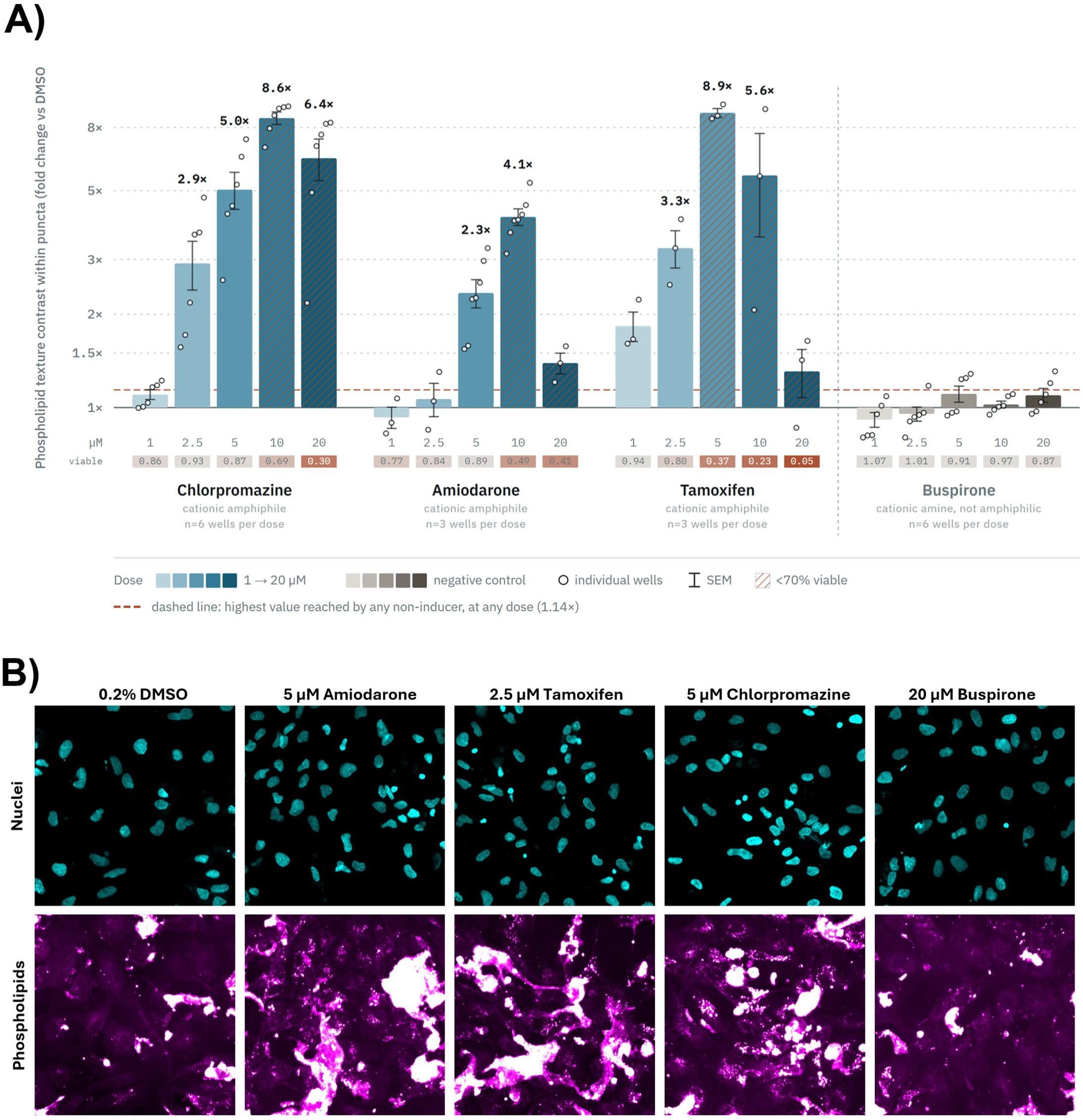
Phospholipid-specific staining confirms phospholipidosis in 92F HLOs treated with cationic amphiphilic drugs. **(A)** Phospholipid puncta texture contrast after 72 h exposure, expressed as fold change over same-plate 0.2% DMSO (log₂ axis). Chlorpromazine, amiodarone, and tamoxifen each increased dose-dependently, reaching 5.04×, 2.34×, and 3.27× while nuclei counts remained at 87%, 89%, and 80% of vehicle, respectively. Buspirone, a lysosomotropic cation that does not induce phospholipidosis, remained flat across the full dose range (0.91–1.10×), and no non-inducer exceeded 1.14× at any dose (dashed line). Bars, mean ± SEM of replicate wells; circles, individual wells. Striped bars indicate doses with fewer than 70% of vehicle nuclei; the strip below the axis gives the viable fraction for each bar. **(B)** Representative images of vehicle- and compound-treated cells. Nuclei, cyan; phospholipids, magenta hot LUT.

## Notes

### Competing Interest Statement

The authors have declared no competing interest.

## References

1. Roadmap to Reducing Animal Testing in Preclinical Safety Studies. https://www.fda.gov/media/186092/download.

2. FDA Announces Plan to Phase Out Animal Testing Requirement for Monoclonal Antibodies and other drugs. U.S. Food and Drug Administration https://www.fda.gov/news-events/press-announcements/fda-announces-plan-phase-out-animal-testing-requirement-monoclonal-antibodies-and-other-drugs (2025).

3. NIH to prioritize human-based research technologies. National Institutes of Health (NIH) https://www.nih.gov/news-events/news-releases/nih-prioritize-human-based-research-technologies (2025).

4. Garcia-Cortes, M. et al. Drug induced liver injury: an update. Arch. Toxicol. 94, 3381–3407 (2020).

5. Reuben, A., Koch, D. G., Lee, W. M. & Acute Liver Failure Study Group. Drug-induced acute liver failure: results of a U.S. multicenter, prospective study. Hepatology 52, 2065–2076 (2010).

6. Hosack, T., Damry, D. & Biswas, S. Drug-induced liver injury: a comprehensive review. Therap. Adv. Gastroenterol. 16, 17562848231163410 (2023).

7. Lin, C. & Khetani, S. R. Advances in engineered liver models for investigating drug-induced liver injury. Biomed Res. Int. 2016, 1829148 (2016).

8. Fisher, J. E. et al. Role of Kupffer cells and toll-like receptor 4 in acetaminophen-induced acute liver failure. J. Surg. Res. 180, 147–155 (2013).

9. Ezhilarasan, D. Hepatotoxic potentials of methotrexate: Understanding the possible toxicological molecular mechanisms. Toxicology 458, 152840 (2021).

10. Ouchi, R. et al. Modeling Steatohepatitis in Humans with Pluripotent Stem Cell-Derived Organoids. Cell Metab. 30, 374–384.e6 (2019).

11. Thompson, W. L. & Takebe, T. Generation of multi-cellular human liver organoids from pluripotent stem cells. Methods Cell Biol. 159, 47–68 (2020).

12. Shinozawa, T. et al. High-Fidelity Drug-Induced Liver Injury Screen Using Human Pluripotent Stem Cell-Derived Organoids. Gastroenterology 160, 831–846.e10 (2021).

13. Zhang, C. J. et al. A human liver organoid screening platform for DILI risk prediction. J. Hepatol. (2023) doi:10.1016/j.jhep.2023.01.019.

14. Meyer, S. R. et al. A high-throughput microphysiological liver chip system to model drug-induced liver injury using human liver organoids. Gastro Hep Adv. 3, 1045–1053 (2024).

15. Rouquié, D. et al. The OASIS Consortium: integrating multi-omics technologies to transform chemical safety assessment. Toxicol. Sci. 208, 225–232 (2025).

16. Ewald, J. D. et al. Cell Painting for cytotoxicity and mode-of-action analysis in primary human hepatocytes. bioRxiv.org (2025) doi:10.1101/2025.01.22.634152.

17. Chalasani, N. et al. Features and Outcomes of 899 Patients With Drug-Induced Liver Injury: The DILIN Prospective Study. Gastroenterology 148, 1340–52.e7 (2015).

18. McCracken, K. W. et al. Modelling human development and disease in pluripotent stem-cell-derived gastric organoids. Nature 516, 400–404 (2014).

19. Zou, W. et al. Sulindac metabolism and synergy with tumor necrosis factor-alpha in a drug-inflammation interaction model of idiosyncratic liver injury. J. Pharmacol. Exp. Ther. 331, 114–121 (2009).

20. Tarazi, E. M., Harter, J. G., Zimmerman, H. J., Ishak, K. G. & Eaton, R. A. Sulindac-associated hepatic injury: analysis of 91 cases reported to the Food and Drug Administration. Gastroenterology 104, 569–574 (1993).

21. Zou, W. et al. Sulindac metabolite and TNFa interaction in liver injury induced by sulindac and lipopolysaccharide in rats. FASEB J. 23, (2009).

22. Begriche, K., Massart, J., Robin, M.-A., Borgne-Sanchez, A. & Fromenty, B. Drug-induced toxicity on mitochondria and lipid metabolism: mechanistic diversity and deleterious consequences for the liver. J. Hepatol. 54, 773–794 (2011).

23. Fromenty, B. & Pessayre, D. Inhibition of mitochondrial beta-oxidation as a mechanism of hepatotoxicity. Pharmacol. Ther. 67, 101–154 (1995).

24. Buchheim, W., Drenckhahn, D. & Lüllmann-Rauch, R. Freeze-fracture studies of cytoplasmic inclusions occurring in experimental lipidosis as induced by amphiphilic cationic drugs. Biochim. Biophys. Acta 575, 71–80 (1979).

25. Tummino, T. A. et al. Drug-induced phospholipidosis confounds drug repurposing for SARS-CoV-2. Science (2021) doi:10.1126/science.abi4708.

26. Halliwell, W. H. Cationic amphiphilic drug-induced phospholipidosis. Toxicol. Pathol. 25, 53–60 (1997).

27. Björnsson, E. et al. Drug-induced autoimmune hepatitis: clinical characteristics and prognosis. Hepatology 51, 2040–2048 (2010).

28. Lewis, W. et al. Fialuridine and its metabolites inhibit DNA polymerase gamma at sites of multiple adjacent analog incorporation, decrease mtDNA abundance, and cause mitochondrial structural defects in cultured hepatoblasts. Proc. Natl. Acad. Sci. U. S. A. 93, 3592–3597 (1996).

29. Bort, R., Ponsoda, X., Jover, R., Gómez-Lechón, M. J. & Castell, J. V. Diclofenac toxicity to hepatocytes: a role for drug metabolism in cell toxicity. J. Pharmacol. Exp. Ther. 288, 65–72 (1999).

30. Williams, D. P., Pirmohamed, M., Naisbitt, D. J., Maggs, J. L. & Park, B. K. Neutrophil cytotoxicity of the chemically reactive metabolite(s) of clozapine: possible role in agranulocytosis. J. Pharmacol. Exp. Ther. 283, 1375–1382 (1997).

31. Dragovic, S., Boerma, J. S., van Bergen, L., Vermeulen, N. P. E. & Commandeur, J. N. M. Role of human glutathione S-transferases in the inactivation of reactive metabolites of clozapine. Chem. Res. Toxicol. 23, 1467–1476 (2010).

32. Proctor, W. R. et al. Utility of spherical human liver microtissues for prediction of clinical drug-induced liver injury. Arch. Toxicol. 91, 2849–2863 (2017).

33. Shah, F. et al. Setting clinical exposure levels of concern for drug-induced liver injury (DILI) using mechanistic in vitro assays. Toxicol. Sci. 147, 500–514 (2015).

34. Schadt, S. et al. Minimizing DILI risk in drug discovery - A screening tool for drug candidates. Toxicol. In Vitro 30, 429–437 (2015).

35. Xu, J. J. et al. Cellular imaging predictions of clinical drug-induced liver injury. Toxicol. Sci. 105, 97–105 (2008).

36. Gerets, H. H. J. et al. Characterization of primary human hepatocytes, HepG2 cells, and HepaRG cells at the mRNA level and CYP activity in response to inducers and their predictivity for the detection of human hepatotoxins. Cell Biol. Toxicol. 28, 69–87 (2012).

37. Sulindac. (National Institute of Diabetes and Digestive and Kidney Diseases, Bethesda, 2012).

38. Weinstein, G. D. Evaluation of possible chronic hepatotoxicity from methotrexate for psoriasis. Arch. Dermatol. 102, 613 (1970).

39. Kremer, J. M. & Lee, J. K. The safety and efficacy of the use of methotrexate in long-term therapy for rheumatoid arthritis. Arthritis Rheum. 29, 822–831 (1986).

40. Whiting-O’Keefe, Q. E., Fye, K. H. & Sack, K. D. Methotrexate and histologic hepatic abnormalities: a meta-analysis. Am. J. Med. 90, 711–716 (1991).

41. Aithal, G. P. Hepatotoxicity related to methotrexate. in Drug-Induced Liver Disease 593–604 (Elsevier, 2013).

42. Chidsey, C. A. et al. Studies of the absorption and removal of propranolol in hypertensive patients during therapy. Circulation 52, 313–318 (1975).

43. Nevirapine. in LiverTox: Clinical and Research Information on Drug-Induced Liver Injury (National Institute of Diabetes and Digestive and Kidney Diseases, Bethesda (MD), 2012).

44. Diclofenac. in LiverTox: Clinical and Research Information on Drug-Induced Liver Injury (National Institute of Diabetes and Digestive and Kidney Diseases, Bethesda (MD), 2012).

45. Dash, A., Figler, R. A., Sanyal, A. J. & Wamhoff, B. R. Drug-induced steatohepatitis. Expert Opin. Drug Metab. Toxicol. 13, 193–204 (2017).

46. Francis, P. & Navarro, V. J. Drug-induced hepatotoxicity. in StatPearls (StatPearls Publishing, Treasure Island (FL), 2025).

47. Okita, K. et al. A more efficient method to generate integration-free human iPS cells. Nat. Methods 8, 409– 412 (2011).

48. Ryu, E. K. et al. Expression and purification of recombinant human fibroblast growth factor receptor in Escherichia coli. Protein Expr. Purif. 49, 15–22 (2006).

49. Björnsson, E. S. & Hoofnagle, J. H. Categorization of drugs implicated in causing liver injury: Critical assessment based on published case reports. Hepatology 63, 590–603 (2016).

50. McKenzie, R. et al. Hepatic failure and lactic acidosis due to fialuridine (FIAU), an investigational nucleoside analogue for chronic hepatitis B. N. Engl. J. Med. 333, 1099–1105 (1995).

51. LiverTox: Clinical and Research Information on Drug-Induced Liver Injury. (National Institute of Diabetes and Digestive and Kidney Diseases, Bethesda (MD), 2012).

52. Olubamiwa, A. O. et al. DILIrank 2.0: An updated and expanded database for drug-induced liver injury risk based on FDA labeling and a literature review. Drug Discov. Today 30, 104485 (2025).

53. Thakkar, S. et al. Drug-induced liver injury severity and toxicity (DILIst): binary classification of 1279 drugs by human hepatotoxicity. Drug Discov. Today 25, 201–208 (2020).

54. Meyer, S. R., Suzuki, M., Jan, K., Hagimoto, M. & Sexton, J. Z. Best practices for cell painting morphologic profiling – every cell in focus. Micros. Today 32, 30–36 (2024).

55. McQuin, C. et al. CellProfiler 3.0: Next-generation image processing for biology. PLoS Biol. 16, e2005970 (2018).

56. Stirling, D. R. et al. CellProfiler 4: improvements in speed, utility and usability. BMC Bioinformatics 22, 433 (2021).

57. Carpenter, A. E. et al. CellProfiler: image analysis software for identifying and quantifying cell phenotypes. Genome Biol. 7, R100 (2006).

58. Serrano, E. et al. Reproducible image-based profiling with Pycytominer. Nat. Methods 22, 677–680 (2025).

