## Supplemental Tables and Methods for "High-Throughput Morphological Profiling of Patient-Derived Human Liver Organoids for Hepatotoxicity Hazard Identification and Compound Prioritization": Supplemental_Methods.docx

Exposure-Anchored Translation of Benchmark Concentrations

### Clinical exposure data: Peak total plasma concentrations (C_max_) were curated for 68 of the 74 screened compounds. Values were taken preferentially from regulatory labels or primary clinical pharmacokinetic studies, and otherwise from published compilations (Xu *et al.* 2008; Walker *et al.* 2020), with each entry annotated by source and confidence. Where independent sources disagreed by more than twofold, the discrepancy and its resolution were recorded. Six compounds were excluded for lack of interpretable human pharmacokinetics: two green-tea catechins (epigallocatechin gallate, gallocatechin), a compound for which the parent and its intravenous ester prodrug could not be distinguished (methyldopa/methyldopate), a peptide without human exposure data (alarelin), and an electrolyte for which C_max_ is not a meaningful quantity (calcium gluconate). For the two fixed-dose combinations in the library, exposure was entered as the molar sum of both components (amoxicillin–clavulanate, 9.03 + 6.32 µM; sulfamethoxazole–trimethoprim, 178 + 5.41 µM).

### Plasma free fractions: The unbound fraction in plasma (*f*_u,p_) was curated for 66 of the 68 compounds with a C_max_. Values were drawn from FDA or EMA product labels where available, otherwise from primary pharmacokinetic literature, and each was assigned a confidence grade. Eleven values were added from regulatory labels or direct equilibrium-dialysis or ultrafiltration measurements; one (azithromycin) was interpolated log-linearly to the clinical C_max_ because its binding is concentration-dependent over the relevant range (51% bound at 0.02 µg/mL to 7% at 2 µg/mL); and three (nicotinamide, 6-thioguanine, floxuridine) were assigned the value of a close structural analogue or drug class in the absence of any human measurement and are flagged as such. Two investigational compounds (IOWH-032, inarigivir) have no published human plasma protein binding and were analysed on a total-C_max_ basis only. All analyses were repeated with the three analogy-based values excluded, and the conclusions were unchanged.

### In vitro dosimetry: The freely dissolved concentration in the culture well was computed for each compound at its own BMC_50_ using the Armitage *in vitro* distribution model (Armitage *et al.* 2014), as implemented in armitage_eval in the httk R package (v2.7.4). The model performs a mass balance across medium water, medium protein and lipid, plate plastic, and cellular storage lipid, membrane lipid and protein, with ionisation handled explicitly via pK_a_.

Assay geometry was set to the conditions of the screen: 384-well format, 50 µL working volume, 8,000 cells per well, and pH 7.4. Culture medium was represented as serum-free hepatocyte medium containing fatty-acid-free bovine serum albumin at 20–50-fold below physiological serum albumin (40 g/L). The geometric midpoint of that range, 1.27 g/L, was used for the primary analysis; because httk scales its serum albumin concentration (24 g/L) by the fetal-bovine-serum fraction, this was implemented as FBSf = 0.0528, which was confirmed to yield 1.26 g/L albumin in the modelled well. Serum lipid was set to zero to reflect the absence of serum. Sensitivity analyses were run at 20-fold and 50-fold below serum albumin (2.00 and 0.80 g/L; FBSf = 0.0833 and 0.0333).

Physicochemical parameters (log K_ow_, aqueous solubility, melting point, Henry’s law constant, and acidic and basic pK_a_) were taken from the curated httk chemical library for 61 of the 74 compounds. The remaining 13 were resolved to PubChem identifiers, and log K_ow_ was computed by the Crippen method and aqueous solubility by the ESOL (Delaney) estimator in RDKit (v2026.03.5); melting point and Henry’s law constant were imputed at the panel median, both of which influence only the solubility-saturation check, which was not limiting at the concentrations tested. Agreement between the two sources was assessed on the 61 compounds present in both and was acceptable for log K_ow_ (Pearson *r* = 0.94, mean absolute error 0.62 log units).

The free fraction in medium was obtained as the ratio of the model’s dissolved concentration to the nominal concentration, *f*_u,med_ = C_dissolved_ / BMC_50_.

As a simpler comparator, a second estimate of *f*_u,med_ was computed from the plasma free fraction alone using the standard protein-dilution relationship, *f*_u,med_ = 1/[1 + *R*(1 − *f*_u,p_)/*f*_u,p_], where *R* is the ratio of medium to plasma albumin (Austin *et al.* 2002; Kilford *et al.* 2008). This accounts for albumin binding only and is reported alongside the full mass-balance result.

### Human-equivalent plasma concentration and offset estimation: Under the free-drug hypothesis, equal freely dissolved concentrations were assumed to drive equal cellular response *in vitro* and *in vivo* (Honda *et al.* 2019). Each compound’s benchmark concentration was therefore expressed as the total plasma concentration that would deliver the same free concentration in a patient:

HEC = BMC_50_ × *f*_u,med_ / *f*_u,p_

The systematic offset between this quantity and clinical exposure was estimated as the geometric mean of HEC/C_max_ across all compounds, with a 95% confidence interval from 5,000 bootstrap resamples of the log_10_ ratios. This yielded an offset of 23-fold (95% CI 12–45), which was interpreted as the aggregate contribution of static culture — the absence of flow and clearance, compound depletion over the 72 h exposure, and the mismatch between sustained *in vitro* treatment and a transient plasma peak. Exposure margins were then computed per compound as

margin = (HEC / offset) / C_max_

and agreement was summarised as the proportion of compounds with |log_10_ margin| below 1 and below 2, corresponding to agreement within 10-fold and 100-fold of clinical C_max_. All summary statistics for concentrations and ratios are geometric means, since both span several orders of magnitude.

This offset is estimated from the same data it is applied to, and is therefore not an independent validation of the central tendency; the quantities that do not depend on it are the width of its confidence interval, the dispersion of the per-compound margins, and the comparison with the independently measured static-to-microphysiological potency shift reported previously for the tenofovir–inarigivir combination. That earlier comparison is nominal-to-nominal and therefore still contains the in-well binding component modelled explicitly here, so the two estimates are related but not interchangeable and were not combined.

### Statistical analysis: Correspondence between translated points of departure and clinical exposure was assessed by Spearman rank correlation. Whether the per-compound pairing was closer than expected by chance was tested by permutation: the assignment of C_max_ to compounds was shuffled 20,000 times and the mean |log_10_ margin| of the observed pairing compared with the resulting null distribution.

Because the dosimetry correction is a function of lipophilicity, and lipophilicity is correlated with clinical exposure, two further controls were used to determine whether any observed correlation was attributable to the organoid measurement rather than to physicochemistry. First, BMC_50_ values were permuted across compounds (3,000 iterations) while the compound-specific correction factor *f*_u,med_/*f*_u,p_ was left intact, and the correlation recomputed. Second, the partial Spearman correlation between the translated point of departure and C_max_ was computed conditional on that correction factor.

Statistical power was assessed by simulation. For each compound, the benchmark concentration that would be observed under perfect compound-specific translation was constructed as BMC_50,ideal_ = C_max_ × offset × *f*_u,p_ / *f*_u,med_. This ideal value was then degraded by the measured experimental limitations: multiplicative log-normal noise with a compound-specific standard deviation derived from the observed inter-donor fold-variability (σ_log10_ = log_10_(fold-variability)/3.53, the expected range-to-SD ratio for *n* = 7; median 0.172 across the panel), followed by censoring to the tested concentration range of 1.23–100 µM. The Spearman correlation and permutation test were then recomputed on the simulated data over 2,000 replicates to establish the correlation that would have been recoverable had genuine compound-specific translation been present.

### Software and data availability: Analyses were performed in Python 3.12 (pandas 3.0.5, SciPy 1.18.0, NumPy 2.4.4, Matplotlib 3.11.1) and R 4.6.1 (httk 2.7.4, data.table 1.18.4), with cheminformatics in Python 3.11 (RDKit 2026.03.5). Analysis scripts, the curated exposure and protein-binding tables with per-value sources, the physicochemical parameter table with provenance for every compound, and the per-compound translation output are provided as supplementary files.

**Confirmation of Drug-Induced Phospholipidosis:** Human liver organoids from iPSC line 92F were exposed to selected compounds for 72 h, followed by staining with Hoechst 33342 (1:1,000; Invitrogen, 62249), and HCS LipidTOX Red Phospholipidosis Detection Reagent (1:1000; Invitrogen, H34351). Images were acquired in the same manner and analyzed using a custom Cell Profiler pipeline to quantify phospholipid-associated puncta and textural features. Phospholipid texture contrast, reflecting the sharpness of segmented phospholipid puncta, was used as the primary phospholipidosis metric and normalized to same-plate 0.2% DMSO controls. Nuclei counts were quantified in parallel as a measure of cytotoxicity.

### Supplemental Methods References:

Armitage JM, Wania F, Arnot JA. Application of mass balance models and the chemical activity concept to facilitate the use of *in vitro* toxicity data for risk assessment. *Environ Sci Technol.* 2014;48(16):9770–9779.

Pearce RG, Setzer RW, Strope CL, Sipes NS, Wambaugh JF. httk: R package for high-throughput toxicokinetics. *J Stat Softw.* 2017;79(4):1–26.

Honda GS, Pearce RG, Pham LL, *et al.* Using the concordance of *in vitro* and *in vivo* data to evaluate extrapolation assumptions. *PLoS One.* 2019;14(5):e0217564.

Austin RP, Barton P, Cockroft SL, Wenlock MC, Riley RJ. The influence of nonspecific microsomal binding on apparent intrinsic clearance, and its prediction from physicochemical properties. *Drug Metab Dispos.* 2002;30(12):1497–1503.

Kilford PJ, Gertz M, Houston JB, Galetin A. Hepatocellular binding of drugs: correction for unbound fraction in hepatocyte incubations using microsomal binding or drug lipophilicity data. *Drug Metab Dispos.* 2008;36(7):1194–1197.

Landrum G. RDKit: open-source cheminformatics. https://www.rdkit.org

Delaney JS. ESOL: estimating aqueous solubility directly from molecular structure. *J Chem Inf Comput Sci.* 2004;44(3):1000–1005.

Paul Friedman K, Gagne M, Loo LH, *et al.* Utility of *in vitro* bioactivity as a lower bound estimate of *in vivo* adverse effect levels and in risk-based prioritization. *Toxicol Sci.* 2020;173(1):202–225.
